# The dependence of forecasts on sampling frequency as a guide to optimizing monitoring in community ecology

**DOI:** 10.1101/2023.06.19.545268

**Authors:** Uriah Daugaard, Stefanie Merkli, Ewa Merz, Francesco Pomati, Owen L. Petchey

## Abstract

Facing climate change and biodiversity loss, it is critical that ecology advances so that processes, such as species interactions and dynamics, can be correctly estimated and skillfully forecasted. As different processes occur on different time scales, the sampling frequency used to record them should intuitively match these scales. Yet, the effect of data sampling frequency on ecological forecasting accuracy is understudied. Using a simple simulated dataset as a baseline and a more complex high-frequency plankton dataset, we tested how different sampling frequencies impacted abundance forecasts of different plankton classes and the estimation of their interactions. We then investigated whether plankton growth rates and body sizes could be used to select the most appropriate sampling frequency. The simple simulated dataset showed that the optimal sampling frequency scaled positively with growth rate. This finding was not repeated in the analyses of the plankton time series, however. There, we found that a reduction in sampling frequency worsened forecasts and led us to both over- and underestimate plankton interactions. This suggests that forecasting can be used to determine the ideal sampling frequency in scientific and monitoring programs. A better study design will improve theoretical understanding of ecology and advance policy measures dealing with current global challenges.

**Open research statement:** Data and code used for the analyses and figures are available on Zenodo: https://doi.org/10.5281/zenodo.10066786. Environmental (lake) data (Merkli et al. 2022) are available from ERIC: https://doi.org/10.25678/00066D.

## Introduction

Ecological processes, such as species abundances and interactions, are temporal scale dependent and can range from short-term responses to perturbations (e.g. Medeiros *et al*. 2023) to long-term adaptations to changed environmental conditions (Crozier and Hutchings 2014). A central aim of ecology is not only to infer such processes from population dynamics, but also to forecast them to advance and test ecological theory and to inform decision-making (Lewis et al. 2023), for example in conservation ecology (Tulloch, Hagger, and Greenville 2020). It is thus intuitively important to record population sizes with sampling frequencies that match the temporal scale of the focal processes. Despite this, little is known about how sampling frequencies affect ecological forecast skill, or how to select appropriate sampling frequencies.

In general, sampling infrequently can yield time series that are too sparse to adequately infer the processes of interest (Estes et al. 2018). Infrequent sampling can cause under- or overestimations (e.g. Queiroz and Ferreira 2009), imprecise estimates (e.g. Taylor and Howes 1994), and non-detection of focal dynamics (Lehtiniemi et al. 2022). Ultimately, this can lead to the non-detection of relations between variables or to spurious associations (Cabella, Meloni, and Martinez 2019), hindering also the reproducibility of results (Estes et al. 2018).

Regarding forecasting, in various scientific fields (e.g. climatology, hydrology, landslide risk management), it has been found that they benefit from high frequency data (e.g. Liu and Han 2013; Leyton and Fritsch 2004; Bozzano, Mazzanti, and Moretto 2018; Arhab and Huang 2023). However, in ecology the few studies that investigated how forecasting is affected by sampling frequency found that sampling more often could both improve and worsen forecasts (Wauchope et al. 2019; Derot, Yajima, and Schmitt 2020). Moreover, to our knowledge neither these studies nor the ones from other scientific fields account for the fact that higher sampling frequencies result in bigger sample sizes (i.e. number of time points), which needs to be controlled for as it can act as a confounding variable by also yielding better forecasts.

While sampling sufficiently often remains a challenge in some systems, in others automated data collection approaches that create high-frequency data have become available (e.g. Kays et al. 2015; Besson et al. 2022). Nevertheless, also in cases like these it is of interest to select appropriate sampling frequencies that avoid unnecessary samplings and associated costs. Yet, sampling frequencies are commonly chosen based on experience and logistics, but remain otherwise often unjustified and potentially unoptimized (Ma, McKindsey, and Johnson 2022).

Generally speaking, the sampling frequency should be high enough that all system-relevant signals are recorded (Isles and Pomati 2021), but not higher. We hypothesize that a candidate sampling frequency for this is the lowest frequency that still yields the highest forecast skill, as this might suggest that all relevant signals have been captured by the data. However, the high-frequency data necessary to test this is often not available and thus alternative criteria to decide the sampling frequency are needed. For instance, in abundances time series the sampling frequency could potentially be selected based on the *per capita* growth rate (henceforth referred to as growth rate) and body size of the focal species. Indeed, species with lower intrinsic growth rates (i.e. longer generation times) are generally larger in size (Gillooly 2000; Bonner 2015), and their abundances tend to be forecasted better (Petchey et al. 2015; Anderson and Gillooly 2020).

In studies where multiple species are present, however, the different growth rates complicate the process of finding the optimal sampling frequency. Further, a higher species richness implies more species interactions (Borrett and Patten 2003) and abundance forecasts depend on how connected the focal species is, with forecasts being better for species that have many but on average weak interactions (Daugaard et al. 2022). Consequentially, the question becomes whether growth rates can guide the selection of sampling frequencies also in multi-species studies. Moreover, species growth rates and the sampling frequency might not only impact the achieved forecast skills, but also the estimation of species interactions, which is a central aim of community ecology but crucially also strongly data-dependent (Marquez et al. 2022). It follows that if both forecasts and interaction estimates are affected by the sampling frequency, relations between them might be undetectable if the sampling design is inadequate.

Here, we aimed at clarifying the influence of sampling design choices on the outcome of quantitative analyses, which ultimately can help the design of experiments and field observations. In a first step, we carried out a simple simulation of abundance time series to test the possibility of selecting the sampling frequency based on species growth rates and to clarify the relation between sampling frequency and abundance forecast skill. We then extended the analysis by using field data and explored whether the results found *in silico* hold in natural systems and estimated the impacts of sampling frequency, number of time points, growth rates and body sizes first on abundance forecasting and in a second part also on the estimation of species interactions. Lastly, we combined the species interaction estimates and abundance forecast skills found in the previous step. We investigated whether the relation between these two quantities found in a controlled laboratory setting (i.e. in Daugaard et al. 2022) is also found in an observational dataset with greater system complexity and whether the sampling design affected this detection.

Alongside the simulations we used a high-frequency lake phyto- and zooplankton dataset. Plankton population dynamics represent an ideal framework for our study. They show variable and fast-paced time series, driven by their short generation times and their interactions with their system components (e.g. climate and nutrients, Philippart et al. 2000; Merz et al. 2023). Plankton is of great importance in marine and freshwater ecosystems because it is at the base of aquatic food webs regulating biomass transfer and element cycles, and their dynamics affect many ecosystem services (Falkowski 2012). There is therefore a strong interest in forecasting plankton dynamics, including algal blooms (e.g. Rousso *et al*. 2020; Woelmer *et al*. 2022).

Using the field data, we separately (1) sub-sampled the time series using different frequencies while keeping the number of time points constant and (2) reduced the number of time points while keeping the sampling frequency constant. We then estimated the interactions between plankton classes and forecasted their abundances. We hypothesized that growth rates could aid the selection of adequate sampling frequencies in the simulations, but that the greater complexity of field data might prevent this. We also anticipated that fewer or sparser samplings would generally result in worse abundance forecasts, and potentially in under- or overestimations of field plankton interactions. Lastly, we also expected that the non-optimization of frequency and number of samplings would result in insufficient statistical power to detect expected associations between variables, such as the one between forecast skill and species interactions.

## Material and Methods

In this study, we first investigated the effects of sampling design choices on forecasts and, second, their effects on species interaction estimates. In the first part we analyzed both simulated and field data, and we used the latter also in the second part. As such, we describe the collection, processing, and sub-sampling of the field data at the beginning of this section before we describe the forecasting methods and analyses.

### Forecast error as a function of sampling design and functional traits

#### Field data collection and processing

We recorded plankton abundance time series in the eutrophic Lake Greifen (northern Switzerland) with an automated plankton monitoring system that uses a darkfield microscope based on the Scripps Plankton Camera system (Orenstein et al. 2020). The camera takes images at two magnifications (5.0x aimed at phytoplankton and ciliates and 0.5x for zooplankton species) every hour for ten minutes with a frame rate of one frame per second. The instrument has previously been calibrated and its performance compared to traditional microscopy methods has been validated (Merz et al. 2021).

For this study, we used images from April 2019 until December 2021 (994 days). We processed these images as described by Merz *et al*. (2021) to extract regions of interest (ROIs, i.e. the imaged individuals). It has been shown that ROIs/sec is a valid proxy of plankton densities and that ROI area is a robust estimate of their body size (Merz et al. 2021). We classified the ROIs with previously trained convolutional neural networks into zooplankton and phytoplankton classes (Kyathanahally et al. 2021; Kyathanahally 2022). For the zooplankton we had the classes Calanoid Copepods, Cyclopoid Copepods, Ciliates, Daphnids, Nauplii and Rotifers. As is commonly done, we grouped the phytoplankton species into six bins of equal width on the log_10_-scale based on their cell size (i.e ROI area). Size-based phytoplankton bins can be studied as a function of environmental conditions (Yvon-Durocher et al. 2010; Marañón 2015). We calculated daily abundances by summing the hourly abundances (ROIs/sec) per size-bin and per zooplankton class (Appendix S1: Figure S3). We further used lake properties available from Merkli et al. (2022), specifically the daily epilimnetic temperature, the mixed layer depth and irradiance and the weekly concentrations of ammonium, nitrate and phosphate. See Appendix S1: Section S1.2 for details regarding the data collection and processing.

#### Processing of recorded time series

The field data included 1.61% missing data for the phytoplankton and 3.72% for the zooplankton groups. We imputed the missing data by using cubic hermite splines to create complete times series with equidistant spaced data points. From the weekly nutrient chemistry data, we imputed daily values using loess (span=0.15). We added noise to all imputed values to avoid statistical artifacts caused by the imputation (Appendix S1: Section S1.2.3). Following commonplace practices (e.g. Benincà *et al*. 2008), we further processed the time series by carrying out a fourth-root power transformation of the data to dampen population spikes and by detrending and standardizing the time series. We detrended the data by regressing the time series against time and henceforth using the standardized residuals as the new time series (Appendix S1: Figure S4).

##### Estimation of maximum net growth rates

Using the untransformed data, we estimated the daily *per capita* growth rates of the plankton classes as *r_T_* = log(*N_T,t_*/*N_T,t_*-τ) /τ, with *N* being the abundance, *T* the target, *t* the time and τ the time step (here τ = 1 day). To avoid overestimations of growth rates caused by noise, we selected the 90^th^ percentile of the positive values of *r_T_* as the maximum net growth rate of each target.

##### Subsampled time series: reducing sampling frequency and number of time points

To have time series of different resolutions we subsampled the daily data using different frequencies as visualized in Figure 1. A constraint was that at the lowest sampling frequency the recorded time series still contained sufficient time points. We set the lowest sampling frequency to 1/12 (i.e. one sampling every 12 days, resulting in 82 time points). So that we could forecast the same time ahead across all sampling frequencies, the other sampling frequencies were: 1/6, 1/3 and 1/1 (i.e. daily sampling). We further constrained all the subsampled time series to also contain 82 time points. To achieve this while also ensuring that we used all time points exactly once per sampling frequency, we created 12 datasets at each sampling frequency (Figure 1). For the daily sampling the 12 datasets were sequential, while for the sampling frequency of once every 12 days their start dates each differed by one day. Accordingly, and as necessary, at the two intermediate sampling frequencies the datasets were both sequential and shifted by one day.

**Figure 1:**
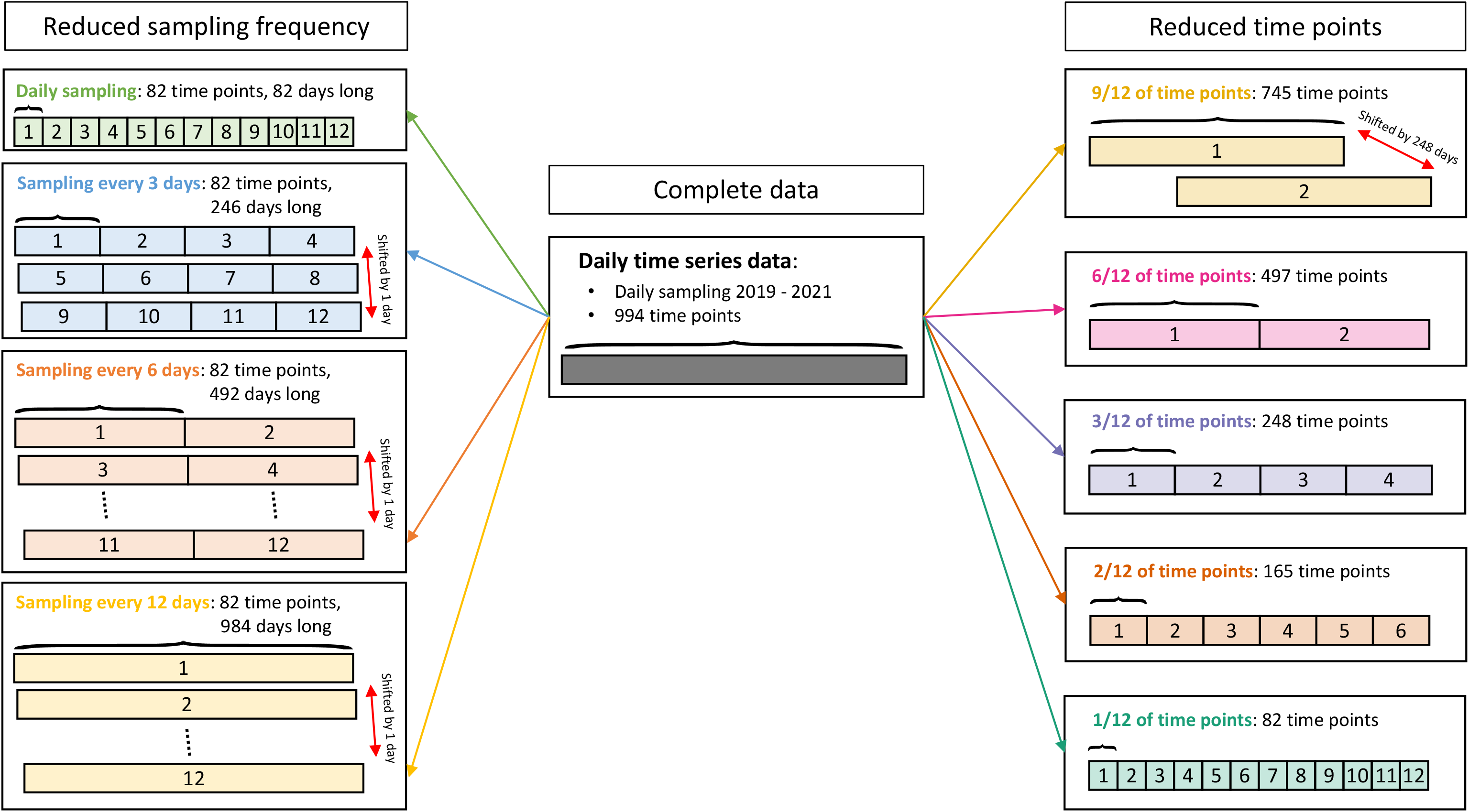
Schematic view of the field datasets used. The middle column shows the complete data. This data is manipulated by reducing the sampling frequency (left column) and by reducing the proportion of kept time points (right column). Each numbered rectangle represents a reduced dataset. In the left column, the red arrows indicate that the sampling start dates of the corresponding time series were shifted by one day so that none of the 12 datasets per sampling frequency used the same time points. In the right column the red arrow indicates by how many days the start date was shifted so that all time points are used at least once.

In summary, with this subsampling method we varied the sampling frequency while controlling the number of data points in the time series. This necessarily resulted in varying absolute time series lengths. For instance, the daily sampled times series are 82 days long, while those with samples every 12 days are 973 days long. This created the possibility that the less frequently sampled times series contained stronger longer-term dynamics (see discussion). In a robustness analysis we controlled for this by keeping the time covered by the time series constant regardless of the sampling frequency (see Appendix S1: Section S2.4.2)

We created time series datasets containing different amounts of time points by keeping the following proportions of the daily data: 9/12, 6/12, 3/12, 2/12 and 1/12 (Figure 1). To use every recorded time point exactly once regardless of the proportion of data retained, we created several datasets at each of the desired lengths (see Figure 1). The exceptions to this are the two datasets that kept 9/12 of the time points as they partially overlapped (Figure 1).

#### Forecasting of species abundances

We forecasted the abundances of taxa from simulated and field data using empirical dynamic modeling (EDM, Ye et al. 2015) using the R-package “rEDM” (Park et al. 2021). In EDM the state of a variable (e.g. the abundance of a taxa) is predicted based on how the variable behaved when it was in a similar state at other times. The similarity of states can be determined using multiple (time-lagged) variables (Takens 1981) and the number of variables used for this is the embedding dimension *E* (Appendix S1: Section S1.3).

##### Forecasting of simulated dynamics

We carried out a simulation study to investigate the relation between sampling frequency, number of time points, growth rate and forecast error. The aim was to have a simple baseline model (i.e. with minimal assumptions and complexity) to inform our research hypotheses under controlled settings and which we then tested in the more complex setting of the field data. We simulated single species time series by using the R-package “odin” (FitzJohn 2022) and the delayed logistic equation (Ruan 2006): 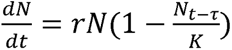. In this equation, the instantaneous rate of change *dN/dt* of the abundance *N* of a taxon depends on the abundance at time point *t* minus τ (the time delay), the growth rate *r* and the carrying capacity *K*. Stable population cycles emerge when *r*τ *>* π*/2*. We varied the number of time points, sampling frequency (between 2 and 1/8 samplings per day), measurement noise and growth rate (between 0.3 and 1.2, with constant *r*τ). We repeated the simulations 100 times for each combination of parameter values.

We used simplex EDM (a single time series technique) with optimized *E* for the forecasts (Sugihara and May 1990). We split the simulated time series into training and evaluation data. We used the former to train the eight days-ahead forecast models (matching the lowest sampling frequency). We then forecasted the abundances in the evaluation data with the models. To assess the forecasts, we calculated the root mean square error (RMSE, i.e. the forecast error). Here, and throughout the study we standardized the RMSE values. The expected value of the standardized RMSE for forecasts based on the average abundance (i.e. the baseline model) is one, which means that forecasts that achieve RMSE values below one perform better than the baseline model (i.e. there is greater forecast skill). See Appendix S1: Sections S1.1 and S1.3.1 for more information.

##### Forecasting of field abundances

We forecasted the abundances of the phyto- and zooplankton classes (henceforth referred to as targets). As predictors we used the targets, the mean epilimnion temperature, the mean mixed layer depth and irradiance, and the concentrations of ammonium, phosphate, and nitrate. We used the EDM technique multiview embedding (Ye and Sugihara 2016). This approach is capable of dealing with high-dimensional systems by fitting all possible low-dimensional (i.e. small *E*) forecast models of which the best ones are then used in an ensemble forecast.

For all the datasets we made 12-days ahead forecasts (matching the lowest sampling frequency) varying the number of steps forecasted ahead based on sampling frequency (Appendix S1: Table S1). With the reduced sampling frequency datasets, we forecasted target abundances using leave-one-out cross-validation (CV). Thus, we refitted the forecasts separately for each time point (i.e., we used the direct forecasting strategy which corresponds to data assimilation at the highest frequency, see Sahoo et al. 2020; Dietze 2017). To reduce computational load, for the complete daily dataset and all other datasets we used *k*-fold CV, by dividing the time series into training data and evaluation data (21 time points, Appendix S1: Table S1). We evaluated forecasts based on the standardized RMSE as a measure of forecast error. We summarized across the *k*-fold CV and across datasets of the same sampling frequency and of the same number of time points by calculating the median RMSE (Appendix S1: Section S1.3.2).

#### Forecast error regressions

We used the calculated forecast errors as the response variable in separate regressions for each of the explanatory variables: (1) sampling frequency; (2) number of time points; (3) target maximum net growth rates; (4) target body size. For regressions (1) and (2) we included an interaction term with a variable indicating whether the targets were zoo- or phytoplankton and we included random intercepts and slopes for each target. For regressions (3) and (4) we used the complete data and log_10_-transformed the explanatory variables to meet the model assumptions.

### Interaction estimates as a function of sampling design and functional traits

#### Estimation of number and strength of interactions

We determined which field targets were causally linked with a test of causation (convergent cross mapping CCM, Sugihara *et al*. 2012). We followed the recommendations of Deyle et al (2016) and extended the methodology with a stringent convergence test that compares CCM skill between two variables using 20% and 50% of the data. This is done with 100 random subsets of the data. If the CCM skill is larger when 50% of the data is used in at least 95% of subsets, then we considered the test to be passed. This convergence test has previously been used by (Merz et al. 2023). In this way we determined the number of interactions of the targets. We did this in every dataset separately.

To estimate the interaction strengths between causally linked state variables we used Smap EDM (Deyle et al. 2016). This method calculates the interaction time series between interacting variables. Similar to the forecasting, this is done by utilizing the information of how the system reacted when it was in a comparable state at other time points. In addition, the nonlinearity parameter *θ* assigns bigger weights to more similar system states. Here we used an extension of this method called regularized Smap EDM (Cenci, Sugihara, and Saavedra 2019), which can handle process noise by introducing a penalization function (i.e. elastic net regularization) and its parameter λ. We determined target-specific values for *θ* and λ with a grid search. More details regarding CCM and Smap are given in Appendix S1: Section S1.4.

To calculate the mean interaction strength of a target, we averaged the absolute values of estimated interaction time series over both time and interacting state variables. To have a single estimate of number and mean strength of interactions per target at each sampling frequency and proportion of time points retained, we averaged over the respective datasets.

#### Interaction estimates regressions

As for the forecast error, we used the number and the average strength of interactions of the targets as response variables in separate regressions with the explanatory variables: (1) sampling frequency; (2) number of time points; (3) target maximum net growth rate; (4) target body size. The used covariates, interactions, random effects and covariate transformations were identical to when the response variable was the forecast error. Models (3) and (4) were Quasi-Poisson regressions for the number of interactions. This was not the case in models (1) and (2) because the number of interactions was not a count but an average calculated over the reduced datasets.

### Effect of sampling design on the detection of correlations between variables

In a final step, we combined the abundance forecasts and the interaction estimates. A recent result shows that forecasts are better for targets with many but on average weak interactions (Daugaard et al. 2022). We tested whether we found this relation also in the field data and whether the sampling frequency and the number of time points influenced the detection. Accordingly, for each sampling frequency and proportion of time points retained we fitted two regressions with the forecast error as the response variable, separately for the two explanatory variables number of interactions and mean interaction strength.

## Results

### Forecast error as a function of sampling design and functional traits

In the simulations, the shape of the relation between sampling frequency and forecast error depended on the growth rate of the target (Figure 2a). The optimal sampling frequency (i.e. the one resulting in the lowest RMSE) increased with target growth rate (Figure 2b). Neither the measurement error nor the number of time points changed this pattern (Appendix S1: Figure S5).

**Figure 2:**
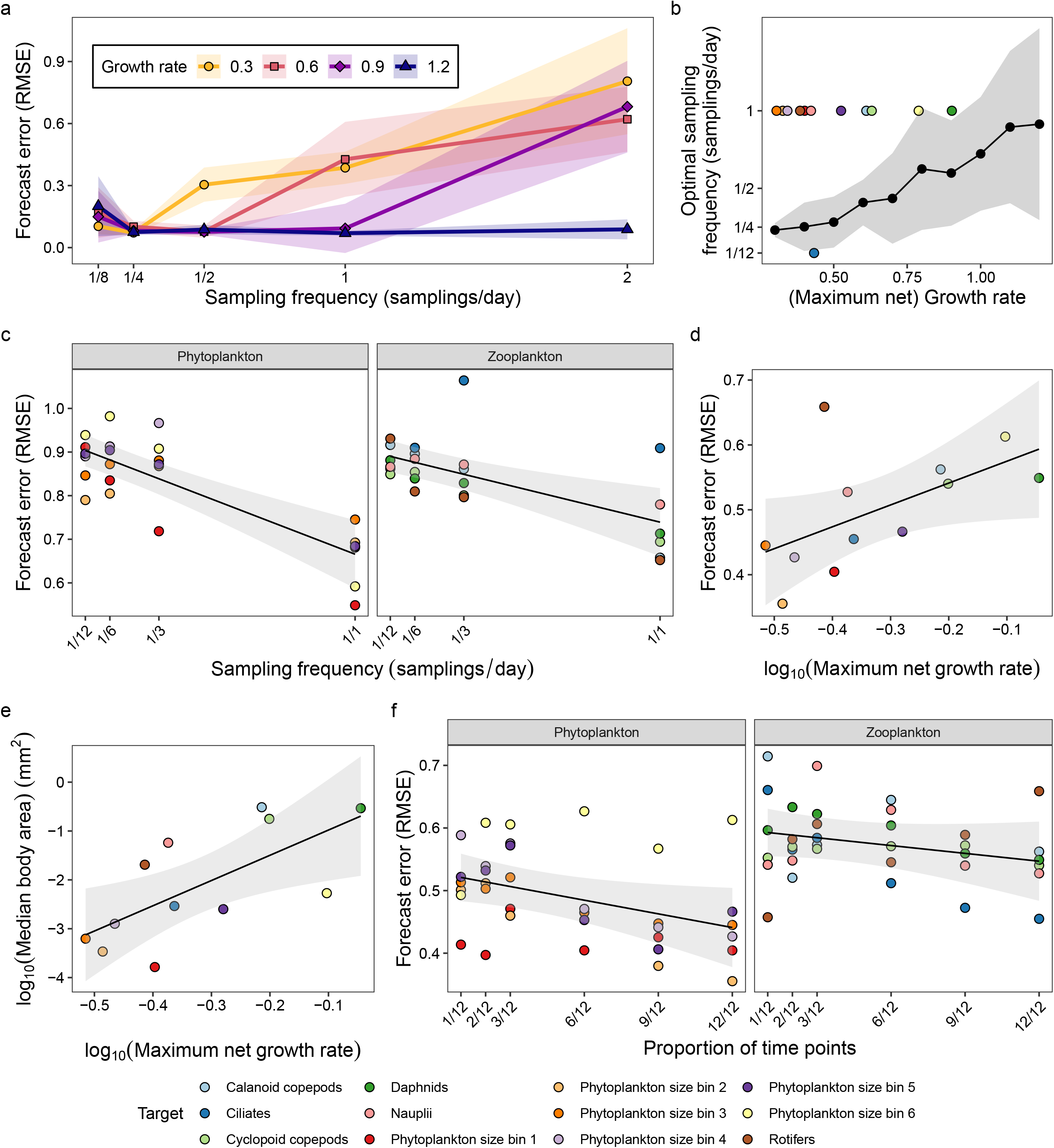
The analyses of forecast error, quantified as RMSE (unitless, as it is based on standardized time series). (a) The relation between sampling frequency and forecast error for the simulated abundance dynamics with different growth rates (75 time points; 60 individuals std. dev. noise; dots: mean forecast errors; shaded region: std. dev. of forecast errors). (b) The relation between growth rate and the optimal sampling frequency for abundance forecasting. Connected black dots (mean values) and the shaded region (standard deviations) show the results for the simulations, with parameter values as in (a). The colored points show the relation for the targets in the field data. (c, d and f) The relations between abundance forecast error (field data) and the (c) sampling frequency, (d) target maximum net growth rate (complete data), and (f) proportion of time points used. (e) The relation between target maximum net growth rates and their median body sizes (complete data). In subfigures (c - f) the dots are colored by target and the lines are the fixed effects of the corresponding regressions (shaded regions: 95% CI).

We found no relation between the maximum net growth rates of the field targets and the optimal sampling frequency, as we achieved the best forecasts with the daily sampling for all but one target (1/12 samplings per day were best for ciliate forecasts, Figure 2b). Across targets, forecast errors decreased linearly by 0.026 and 0.016 for every increase of 0.1 samplings per day (*F*_1,10_ = 42.91, *p* < 0.0001, Figure 2c, Appendix S1: Table S2), respectively for the phyto- and zooplankton targets with no difference between the two groups (*F*_1,10_ = 2.15, *p* = 0.1734). These results were independent of the chosen forecast method as they did not qualitatively change when we used Random Forest to forecast the abundances instead of EDM (Appendix S1: Section S2.4.1). Further, we also found the same relations when we kept the time window covered by the time series constant regardless of the sampling frequency (Appendix S1: Section S2.4.2). Lastly, we found consistent results (i.e., a negative trend between sampling frequency and forecast error) when we forecasted 24 days ahead instead of 12 days (Appendix S1: Section S2.4.3).

We found weak evidence in the complete data that the forecast error increased by 0.337 for every tenfold increase in target maximum net growth rate (*t*_10_ = 2.22, *p* = 0.0509, Figure 2d, Appendix S1: Table S2). Targets with larger body sizes showed bigger maximum net growth rates (log_10_-log_10_ scale, slope = 5.168, *t*_10_ = 2.94, *p* = 0.0147, Figure 2e, Appendix S1: Table S3). Target body size and forecast error were positively correlated (Appendix S1: Figure S6).

Across targets, increasing the proportion of retained time points resulted in lower forecast errors (phytoplankton slope: −0.087; zooplankton slope: −0.050; *F*_1,10_ = 5.67, *p* = 0.0386, Figure 2f, Appendix S1: Table S2). While there was no significant difference in slope between the two plankton groups (*F*_1,10_ = 0.418, *p = 0.5323*), the forecast errors were lower for the phytoplankton targets (intercept difference: −0.069, *F*_1,10_ = 5.20, *p* = 0.0457).

### Interaction estimates as a function of sampling design and functional traits

Increasing the sampling frequency by 0.1 samplings per day increased the estimated number of interactions by 0.225 across the phytoplankton targets (*t*_10.8_ = 2.76, *p* = 0.0190, Figure 3a, Appendix S1: Table S4), but not for the zooplankton targets (*t*_10.8_ = 0.03, *p* = 0.9793). The sampling frequency had no effect on the estimated mean interaction strengths of the targets (*F*_1,10_ = 0.13, *p* = 0.7245, Figure 3b, Appendix S1: Table S5), regardless of the plankton group.

**Figure 3:**
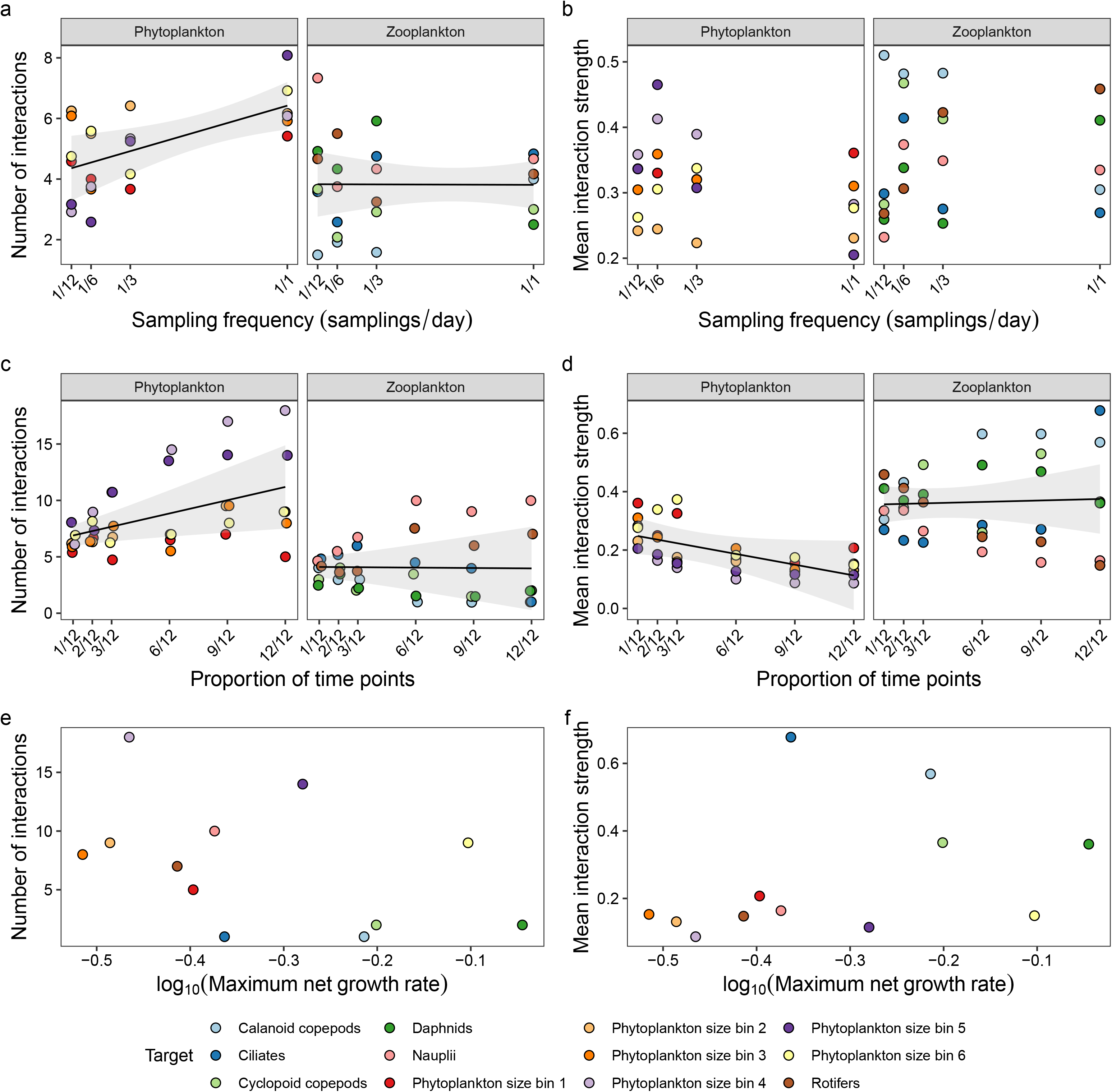
The interaction analyses. (a - b) The effect of sampling frequency on the estimated number of interactions (a) and the mean interaction strengths (b). (c - d) The effect of the proportion of time points used on the number of interactions (c) and the mean interaction strengths (d). (e - f) the relation between the target maximum net growth rate (complete data) and the number of interactions (e) and the mean interaction strengths (f). The dots are colored by target and in (c) they are jittered horizontally to avoid overplotting. The lines are the fixed effects of the corresponding regressions (shaded regions: 95% CI), if a significant relation was found.

For the phytoplankton, decreasing the proportion of time points by 10% decreased the estimated number of interactions by 0.469 (*t*_10_ = 2.69, *p* = 0.0227, Figure 3c, Appendix S1: Table S4) and there was weak evidence that it increased the estimated mean interaction strengths by 0.015 (*t*_10_ = 1.85, *p* = 0.0943, Figure 3d, Appendix S1: Table S5). Across the zooplankton targets, the proportion of time points used did not affect the interactions estimates (*t*_10_ = 0.08, p = 0.9376 and *t*_10_ = 0.25, *p* = 0.8069, respectively for the number and the mean strength of interactions), but we detected target-specific trends (Appendix S1: Figure S7).

The maximum net growth rates were unrelated with the number of interactions (*t*_10_ = 1.41, *p* = 0.1890, Figure 3e, Appendix S1: Table S4) and their mean strength (*t*_10_ = 1.17, *p* = 0.2686, Figure 3f, Appendix S1: Table S5), as was the body size (Appendix S1: Figure S6).

### Effect of sampling design on the detection of correlations between variables

We found no relation between the interaction estimates and the forecast error in the reduced sampling frequency data (Figures 4a and 4b, Appendix S1: Table S6). With the data containing 9/12 of the time points we found a negative relation between the number of interactions and the abundance forecast error (Figure 4c). With the data containing 9/12 and 6/12 of the time points we found a positive relation between the mean interaction strength and the abundance forecast error (Figure 4d). For the complete data and the other data containing a reduced number of time points the estimated slopes between the interaction estimates and the abundance forecast errors also matched in sign with the results of Daugaard *et al*. (2022, Figures 4e and 4f), but with confidence intervals overlapping zero (Figures 4c and 4d; exception: data containing 1/12 of the points in Figure 4d). We further confirmed these relations with the complete data when we made one day ahead forecasts, as done by Daugaard *et al*. (2022, Appendix S1: Figure S10). The number and mean strength of interactions were negatively correlated (Appendix S1: Figure S9).

**Figure 4:**
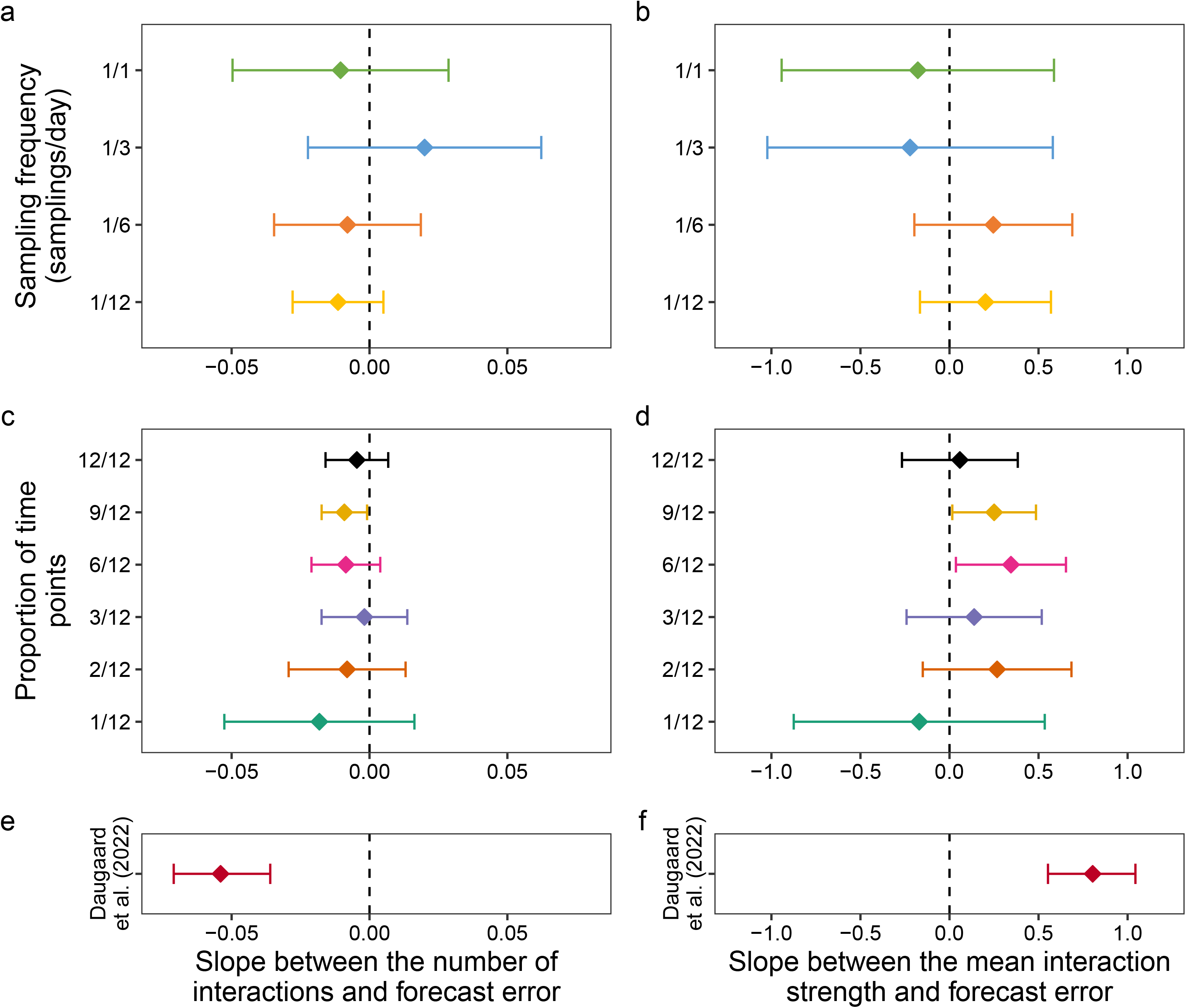
Estimated slopes between the number of interactions of the targets and their abundance forecast error (left column) and between their mean interaction strength and forecast error (right column). (a - b) Reduced sampling frequency analysis. (c - d) Reduced time points analysis. (e - f) Results reported in Daugaard et al. (2022, constant temperature case). The error-bars denote 95% CIs. The datasets differed in the number of targets (i.e. regression sample size): Lake Greifen data had 12 targets, and Daugaard et al. (2022) had eight targets with nine replicates.

## Discussion

We show that abundance forecasts are negatively affected by a reduction in sampling frequency across almost all targets. Further, our forecasts were better for targets with smaller maximal net growth rates and for targets with smaller body sizes. Despite this, we found that growth rates are useful indicators of the optimal sampling frequency only in the simulated single-species time series, but not in the field data. The estimation of interactions also depended on sampling design, as we estimated the phytoplankton targets to interact more the denser the time series were.

Sampling frequency has widely been recognized as having significant impacts on various analyses (e.g. Lehtiniemi et al. 2022; Ma, McKindsey, and Johnson 2022), and yet sampling frequencies are commonly too low (Estes et al. 2018). The implications for ecological forecasting are not well known, as we are aware of few and contrasting findings regarding the effects of sampling frequency on forecasting (Wauchope et al. 2019; Derot, Yajima, and Schmitt 2020), and which potentially were influenced by sample sizes. We found that lowering the sampling frequency worsened the abundance forecasts for 11 out of the 12 targets. Because forecasting is one of the central aims of ecology (Dietze et al. 2018), its dependence on sampling frequency exemplifies the importance of thoughtful sampling design. Failure to carefully calibrate the sampling frequency to match the process of interest could result in inadequate estimations and forecasts, which ultimately could lead to wrong conclusions and inappropriate policies. Notably, the choice of the sampling frequency is part of a larger set of decisions that need to be made so that skillful forecasts can be achieved. This set includes, for example, the choice of the forecasting method, of the data assimilation strategy, of the predictors and of the validation and benchmarking approaches (see Dietze 2017).

Conversely, because forecasts can be evaluated quantitatively their dependence on sampling frequency shows their potential to guide sampling designs. Skillful forecasts are not only an objective in ecology, but they can also advance its theoretical understanding (Lewis et al. 2023). Building on this, we argue that besides improving theory and informing decision-makers, forecasting can guide the design of scientific studies and monitoring programs. Adjusting the sampling frequency based on forecast skill is likely to improve the time series so that the process relevant signals are captured, which is crucial in community ecology (Isles and Pomati 2021).

We found that we estimated fewer interactions for the phytoplankton targets when we decreased the frequency of the samplings. As we achieved the best forecasts at the highest sampling frequency, we expect the estimated number of interactions to be most accurate at this frequency and underestimated otherwise. While the true target interactions are unknown and thus the correctness of the estimations cannot be evaluated, this may become possible thanks to recent advances in the manipulation of species interactions in microcosm experiments (Hu et al. 2022).

The high-frequency data needed to select the sampling design based on forecasts is not always available. In such cases, alternative criteria to decide the sampling frequency are needed. We found that selecting based on growth rates was possible in simulated single-species dynamics but not in the field data, although we achieved better forecasts for targets with smaller growth rates (which also had smaller body sizes, see discussion in Appendix S1: Section S2.1). Thus, in natural systems more frequent sampling is needed than what would be expected based on growth rates. While this could be because of a mismatch between the true target division rates and the estimated maximum net growth rates, the found relation between the latter and the abundance forecasts makes this less likely. Instead, we hypothesize that the reason is the greater complexity of the natural system. Indeed, the Nyquist-Shannon sampling theorem used in signal processing theory states that to correctly record a time series the sampling frequency needs to be at least twice as high as the highest frequency present in the true time series (Shannon 1949). With interacting variables leaving imprints in each other’s time series (Takens 1981), to adequately capture the dynamics of a target it may be necessary to select the sampling frequency not based on the target itself, but on the interacting variable with the fastest time series (e.g. the species with the highest growth rate).

The need for high-frequency data is further corroborated by our result that the effects of sampling design on forecasts and interaction estimations were compounded. We were not able to reproduce the laboratory result that the forecasting of a target depends on the number and strength of its interactions (Daugaard et al. 2022), unless we used the high-frequency field data containing enough time points. Auspiciously, the need for high-frequency data in ecology is increasingly being covered (e.g. Pomati et al. 2011; Besson et al. 2022).

Reducing the number of time points worsened forecasts and, in the case of the phytoplankton groups, resulted in fewer interactions being estimated as significant and thus likely in an underestimation of their actual number. Contrasting this, for the zooplankton groups the interaction estimates were, on average, unaffected by both the used number of time points and the used sampling frequency. A potential explanation for these differences between phyto- and zooplankton might be that the higher trophic level (or other trait differences) of the zooplankton groups renders the estimation of their interactions less data demanding. However, it is also possible that these differences have arisen because of the necessarily different grouping approach (i.e. taxonomically vs. morphologically). However, a closer look at the relation between sample size and interaction estimates shows that the decrease in sample size led to target-specific over- and underestimations of interactions for both the zoo- and the phytoplankton groups (Appendix S1: Figure S7). This suggests that the found differences are not caused by differences between the zoo- and phytoplankton groups but by other differences among targets. Moreover, the fewer time points we used, the more the interaction estimates became similar across the targets (Appendix S1: Figure S8), indicating a loss in power to correctly estimate them. This further shows the importance of adequate sampling design for quantitative analyses.

Varying the sampling frequency necessarily results in time series that either differ in their time range or in number of time points. We controlled for the latter as we considered it to be a stronger confounder and attempted to control for different time ranges with our sub-sampling and forecasting approaches. Yet, it is possible that the different time ranges influenced the results, for instance because the species network may have changed over time (Merz et al. 2023). However, because we only considered a relatively short time and used grouped data which is less likely to change, we do not expect this to be the case. Nevertheless, as a robustness analysis we repeated the forecasting with fixed time ranges across sampling frequencies and confirmed our results (Appendix S1: Section S2.4.2).

In conclusion, we clarify the role of sampling frequency in ecology for the forecasting and estimation of processes. As we face climate change and biodiversity loss (Bellard et al. 2012; Cardinale et al. 2012), ecological forecasting is a field of increasing importance (Dietze 2017). Our results have the potential to improve the design of experiments and field observations, and the estimation of species interactions (an important aspect of biodiversity). Especially the use of forecasts not only as an aim but also as a tool shows promise in this regard. Ultimately, better study designs will improve ecological inference and forecasting, which is fundamental for a better theoretical understanding of ecology and for the implementation of better performing policies and measures that deal with current global challenges.

## Author contributions

SM, EM & FP collected, curated and processed the data. UD & OLP conceived the research questions and designed the analyses. UD analyzed the data and wrote the manuscript. All authors contributed critically to the drafts and gave final approval for publication.

## Conflict of interest statement

The authors declare no conflicts of interest.

## Notes

### Competing Interest Statement

The authors have declared no competing interest.

### Summary of Updates

Improved manuscript with new robustness analyses and implemented revisions.

