## Appendix S1 for "The dependence of forecasts on sampling frequency as a guide to optimizing monitoring in community ecology"

**Journal**: Ecosphere

**Authors**: Uriah Daugaard, Stefanie Merkli, Ewa Merz, Francesco Pomati, Owen Petchey

**Appendix S1**

### **Section S1: Extended methods**

#### **Section S1.1: Simulation Study**

We simulated single species time series by using the R-package “odin” (FitzJohn 2022) and the delayed logistic equation (Ruan 2006): $\frac{dN}{dt}=rN(1-\frac{N_{t-\tau}}{K})$. In this equation, the instantaneous rate of change $dN/dt$ of the abundance $N$ of a species depends on the species’ abundance at time point $t$ minus $\tau$ (the time delay), its growth rate $r$ and its carrying capacity $K$. Stable population cycles emerge when $r\tau>\pi/2$.

As we were interested in the effect of growth rate, we kept the product $r\tau$ constant at 2.3 but used growth rates ranging from 0.3 to 1.2 (which reduces the period of the cycles but does not otherwise change the dynamics, see Figure S1), with a 0.1 step. As the carrying capacity we used $K=500$ individuals (note that the carrying capacity did not affect the results because we standardized the forecast error, see Section S1.3.1). After we simulated the dynamics (with $dt=0.01$ days) we removed the initial transients and added measurement noise with a normal distribution (truncated to avoid negative abundances) to create more realistic time series. We used different levels of noise (standard deviations of 30, 60 and 90 individuals) and for each we repeated the simulations 100 times. Afterwards, we sub-sampled the simulated time series using the following intervals: 0.5, 1, 2, 4 and 8 days (Figure S2). We constrained the time series to have the same number of time points across all sampling intervals. Lastly, we repeated the whole simulation using different time series lengths (number of time points: 45, 60, 75, 90 and 105 days).

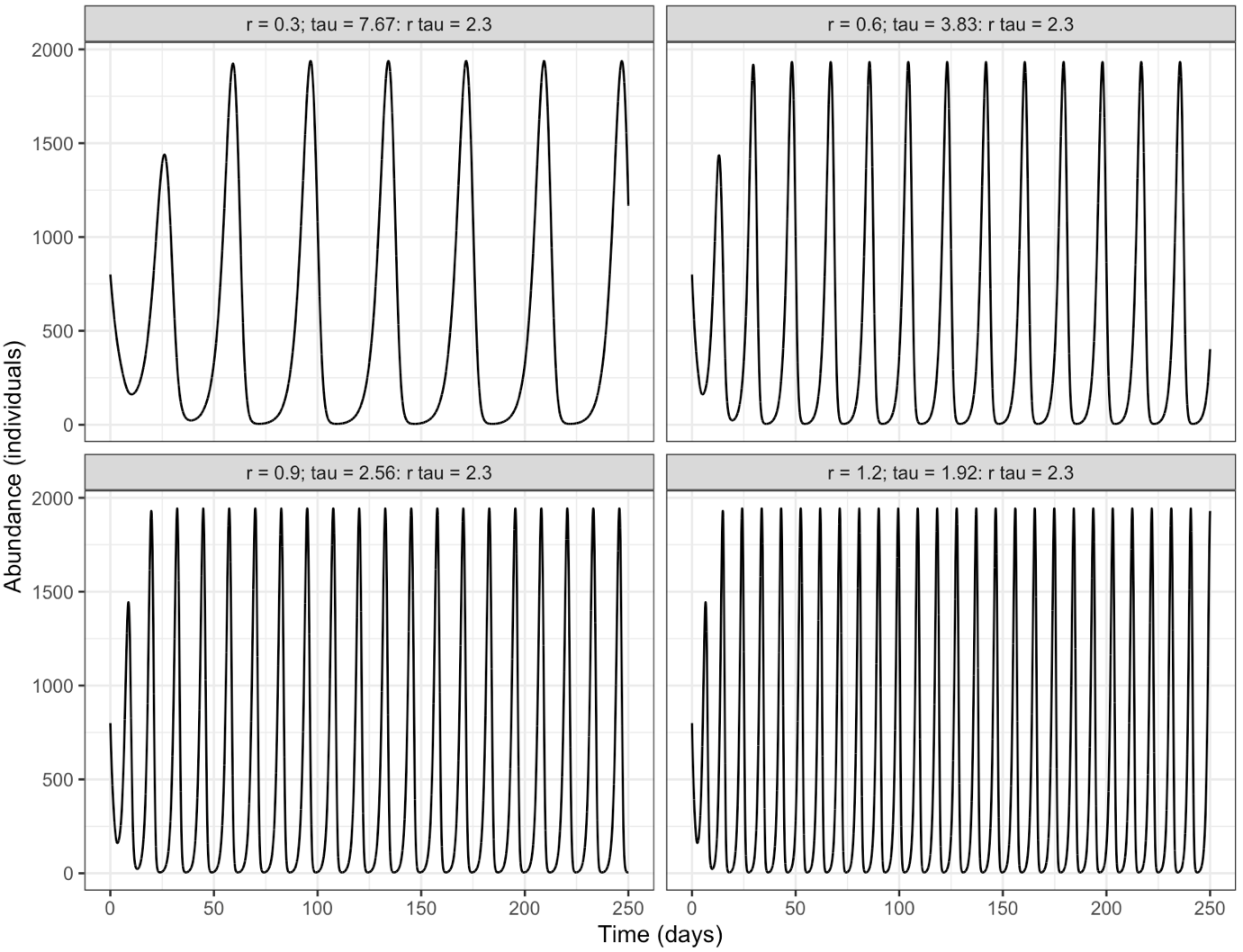

**Figure S1**: Increasing the growth rate in the simulations but keeping the product $r\tau$ constant yields dynamics that show faster cycles. The subpanels show the dynamics for different values of the growth rate, as specified.

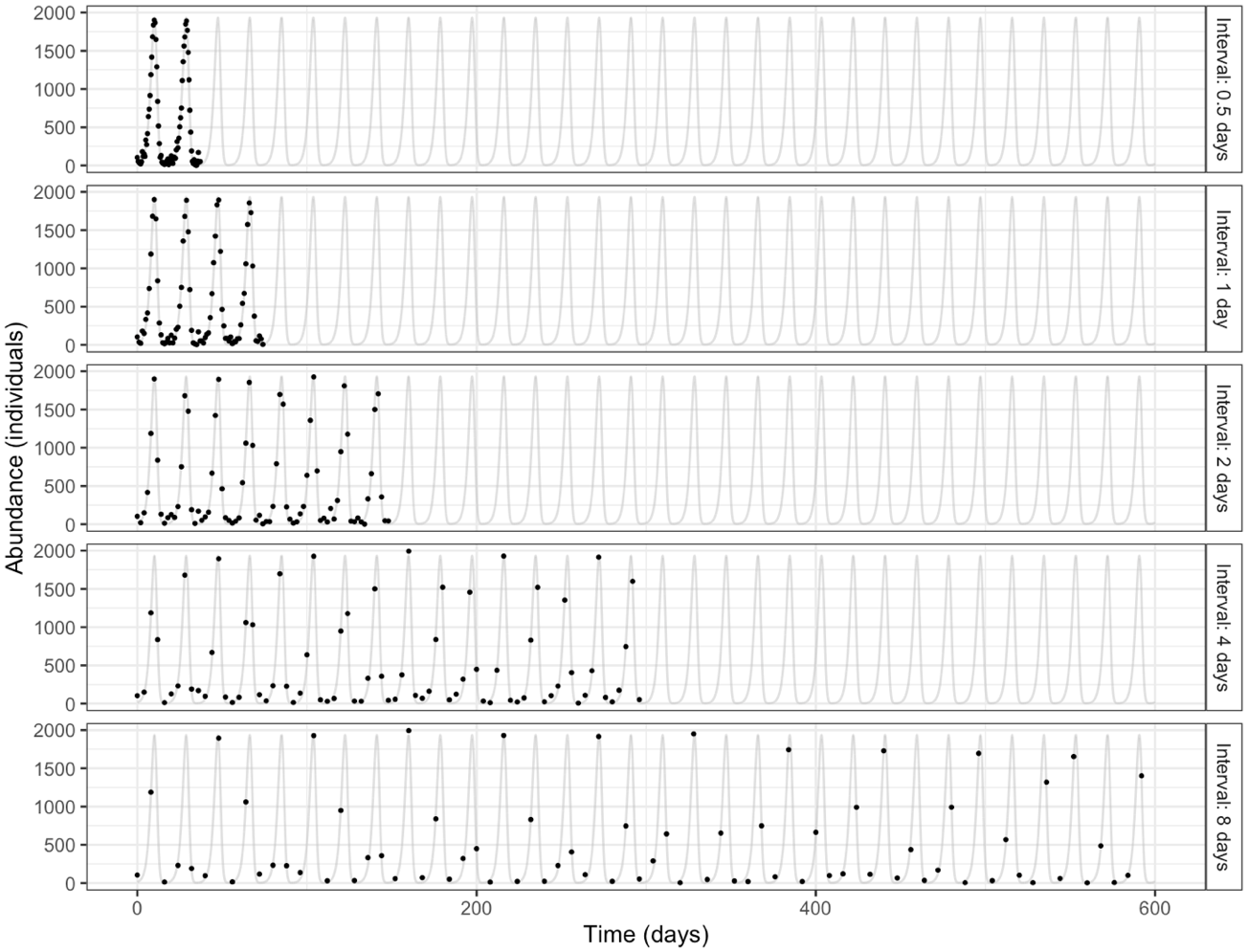

**Figure S2**: Example figure that shows the sub-sampling in the simulated data. the subpanels show the sampled time series with different sampling intervals (i.e. sampling frequencies) as dots. The gray lines show the true time series (without noise). For this figure the parametrization was as follows: r = 0.6; number of time points = 75; standard deviation of added noise = 60 ind.

#### **Section S1.2: High frequency field data**

##### **Section S1.2.1: Data collection**

Lake Greifen is a eutrophic lake in the north of Switzerland with a maximal depth of 32 m and a total area of 8.45 km^2^. We acquired the data at the Eawag automated monitoring station at the northern end of the lake (47.35°N, 8.68°E). We tracked plankton at three meter depth with a dual-magnification darkfield microscope, based on the Scripps Plankton Camera system (the DSPC, Orenstein *et al.* 2020), installed at the monitoring station ([www.aquascope.ch](http://www.aquascope.ch)). The DSPC features a 0.5 times magnification (0p5x), targeting zooplankton and large colony-forming phytoplankton, and a 5.0 times magnification (5p0x), targeting mostly phytoplankton and ciliates. For details about the instrument’s calibration and performance, see Merz *et al.* (2021). We imaged plankton every hour for ten minutes with a frame rate of one frame/second.

We automatically tracked lake physical and chemical variables, including temperature, photosynthetically active radiation (PAR) and oxygen concentration four to eight times a day using a CTD-probe (Ocean Seven 316) and a profiler (<https://www.idronaut.it/>). Once per week, in the same location we took water samples from 3 m depth near the DSPC and analyzed them for nutrient chemistry.

###### **Phytoplankton**

We converted raw regions of interest (ROIs) from the 5p0x magnification of the DSPC into statistics (features files) and image mosaics. We processed the images through color conversion, edge-detection, segmentation, morphological feature extraction, foreground masking and inverse filtering of masked foreground as described elsewhere (Merz *et al.* 2021). Previous work comparing the number of ROIs/sec with traditional microscopy methods has shown that the ROI/sec metric is a valid proxy for phytoplankton density (number of individuals / L), and also that the body size estimates (area of ROI) significantly and robustly scaled with the same measurements derived by traditional microscopy (Merz *et al.* 2021). We classified the obtained ROIs with a previously trained convolutional neural network (CNN) classifier (<https://github.com/kspruthviraj/Plankiformer>) in order to remove images of zooplankton, unidentified entities and dirt from the size distribution of phytoplankton (i.e. area of the imaged plankton objects). From the remaining phytoplankton ROIs, we removed the first and last percentile of the log_10_ distribution of ROI-areas (henceforth referred to as size). We divided the total 98% range of log_10_-transformed size values into six bins of equal width, across the entire dataset, excluding the extreme 2% of data (see Section S1.2.2 for the justification and motivation of the size-binning). We then added the first percentile to the first bin and the last percentile to the last bin. Note that individual phytoplankton cells or colonies are assigned to different bins irrespective of their taxonomic identity, and solely based on each individual object size. This is due to taxa spanning a range of sizes, as well as partially cropped images which lead to different sizes of the same taxon. Finally, we summed the abundances (ROI / sec) per bin per day.

###### **Zooplankton**

For the zooplankton abundances, we subsampled our data from the 0p5x magnification by extracting one frame every 6 seconds to minimize repeated imaging of the same object (Merz *et al.* 2021). We then classified each image using a previously trained CNN (Kyathanahally *et al.* 2021). We had a specific CNN trained for the ciliates in this study ([www.github.com/mbaityje/plankifier/releases/tag/v1.2.2](http://www.github.com/mbaityje/plankifier/releases/tag/v1.2.2)), and used it to classify these organisms from the 5p0x magnification data. We calculated the daily mean abundance (ROI / sec) of each of the zooplankton classes.

###### **CTD data**

Here we describe how we measured and estimated lake properties. Note that we did not use all of them in the final version of this study (see main text).

From the CTD probe, we extracted the average value of each variable across the photic zone (0-8m) for each profile. We then averaged those values across the day.

As measures of water column physical structure, we calculated thermocline depth, mixed layer depth, epilimnetic temperature, and Schmidt stability using the R package rLakeAnalyzer (Winslow *et al.* 2019) and CTD data from the monitoring station (Pomati *et al.* 2011). For the estimation of all variables, we used the temperature profiles generated by the CTD-probe. For the estimation of epilimnetic temperature and Schmidt stability we used lake topology (depth and the lake area at each depth) in addition to the temperature profiles. If no thermocline was detected using a minimum density gradient of 0.1, we used site depth as thermocline depth. We calculated daily averaged values for all variables described above. We further calculated the depth of the oxycline. For that, we set the oxycline depth to the depth where an oxygen threshold of 4 mg / L was reached (a lower threshold was not possible since data were not available from the bottom of the lake; 4 mg / L level allowed us to track relative changes in oxygen across the whole seasonal progression, which we would miss with a lower threshold). If the threshold was not reached at any depth, we used the site depth as oxycline depth. As a measure of available light, we calculated mixed layer irradiance from PAR profiles: we used photosynthetically active radiation (PAR) profiles by the CTD probe at midday (12:00 or 13:00) and excluded the top 3 meters (the monitoring platform shades the CTD-probe). We calculated the light extinction coefficient $K_{e}$ as the slope of the log-transformed PAR against depth regression line. Then, we calculated mixed layer irradiance $I_{mix}$ from $K_{e}$ and mixing depth according to the following equation:

$$I_{mix}=I_{0}\frac{1-e^{-K_{e}z_{mix}}}{K_{e}z_{mix}},$$

Where $I_{mix}$ = mean mixed layer irradiance, $I_{0}$ = irradiance at 3m depth, and $z_{mix}$ = mixed layer depth (or maximal depth if the site was unstratified).

##### **Section S1.2.2: Motivation and justification of phytoplankton binning**

Morphological characteristics of plankton such as individual volume, length, surface and coloniality scale with species functional properties, such as growth rate, sinking rate, grazing resistance, and therefore with population abundances and responses to both abiotic and biotic environmental gradients (Andersen *et al.* 2016; Litchman & Klausmeier 2008). Cell size in particular links the principal axes of resource competition and trophic interactions in plankton (Acevedo-Trejos *et al.* 2015; Marañón 2015). In phytoplankton, large cells have low surface / volume ratio and are less efficient than small cells in nutrient uptake and are therefore at a disadvantage under low nutrient conditions. In contrast, large cells or colonies are less susceptible to being grazed than small cells, for example due to gape limitations in zooplankton. The relationship between size and abundance is an essential link between the individual- and population-level responses to the environment, regulation of community structure, resource allocation among organisms, and the dynamics of food-webs (Barton *et al.* 2013; Marañón 2015; White *et al.* 2007). A common way of grouping plankton into size classes, and studying the dynamics of this important trait, is to report the abundance distribution of taxa as a function of body length or volume in a log–log space (Sprules & Barth 2016). This is often performed by logarithmic binning of organism body-size in the entire local community, and each bin’s abundance (or the scaling of abundances with size) can be studied as a function of environmental conditions (Marañón 2015; Pomati *et al.* 2020; Sprules & Barth 2016; Yvon-Durocher *et al.* 2010).

##### **Section S1.2.3: Further processing of recorded time series**

Figure S3 shows the time series of the plankton groups before the processing described in this section, and Figure S4 displays them after the it.

For the daily time series there was 1.61% missing data for the phytoplankton and ciliates and 3.72% missing data in the zooplankton groups. We imputed the missing data by using cubic hermite splines to create complete times series with equidistant spaced data points. To avoid statistical artifacts caused by the imputation, we added noise to the imputed values as follows. For a given imputed time point we first used a sliding window approach that included the previous and successive 20 time points to calculate the variance of the temporally detrended time series. We then used this variance in a normal distribution (truncated to avoid negative abundances) to randomly draw the noise added to the imputed point. The exception to this were the concentrations of nitrate, phosphate and ammonium that were measured only once per week. We imputed these time series using loess (span=0.15) and then used the variance of the resulting residuals as before to add noise to the imputation.

Following commonplace practices (e.g. Benincà *et al.* 2008), we further processed the time series by carrying out a fourth-root power transformation of the data to dampen population spikes and by detrending and standardizing the time series. We detrended the data by regressing the time series against time and henceforth using the resulting standardized residuals as the new time series.

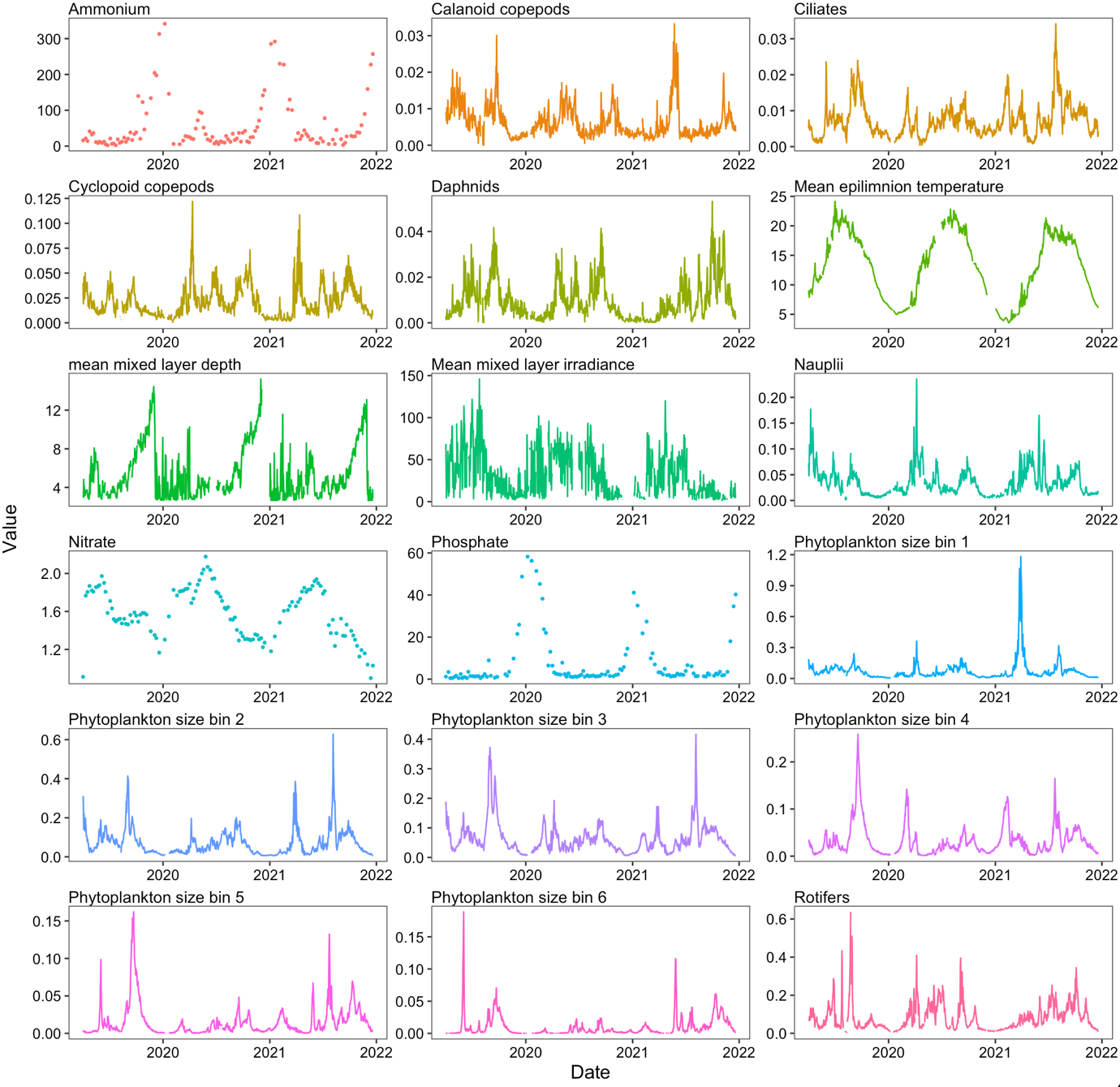

**Figure S3**: The recorded dynamics of the variables used. For the plankton groups the y-axis is in ROI / sec, for ammonium and phosphate it is in µg/L, for nitrate it is in mg/L, for the mean epilimnion temperature it is in degrees Celsius, for the mean mixed layer depth in meters and for the mean mixed layer irradiance it is in Wh m^-2^ d^-1^.

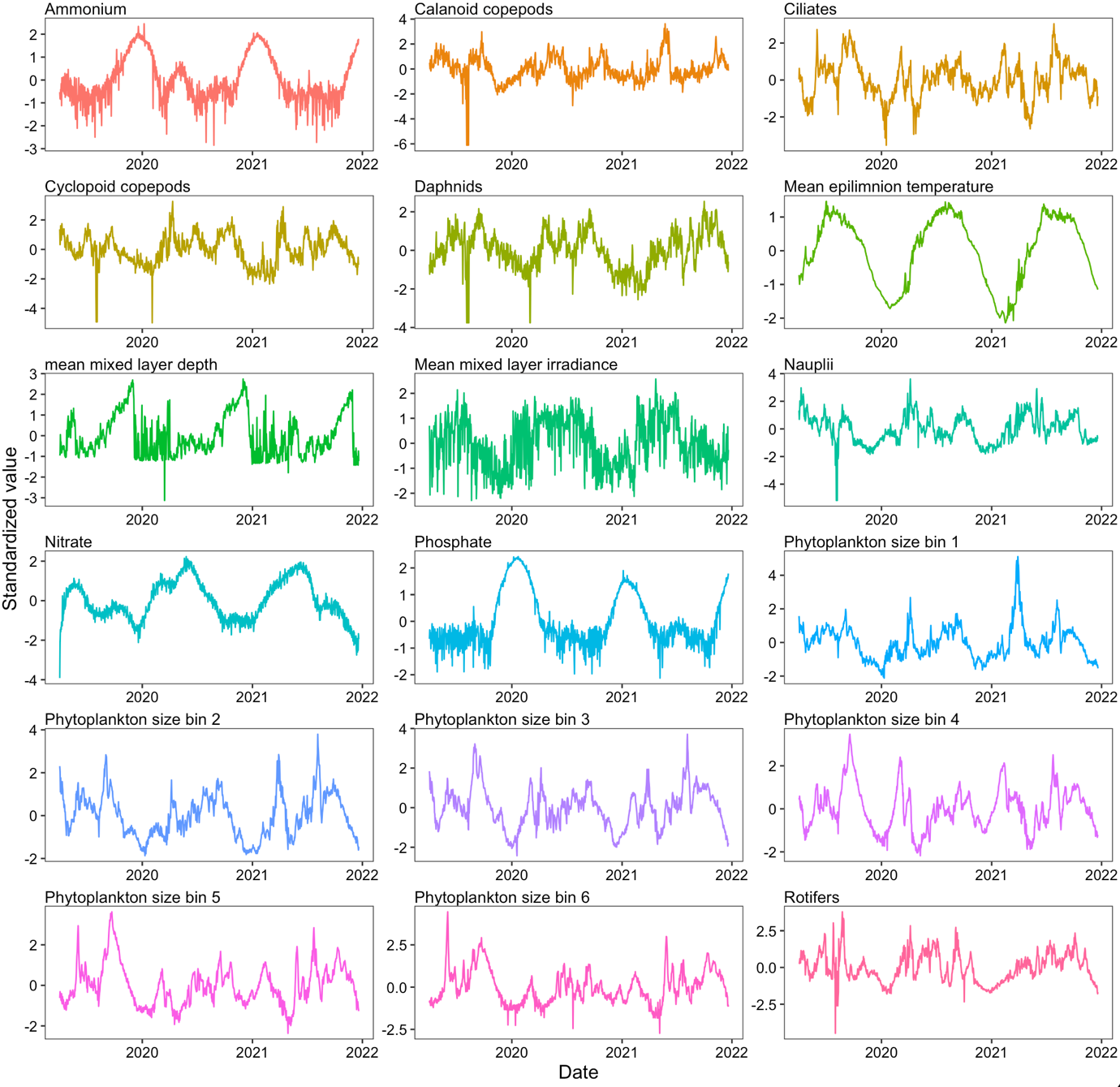

**Figure S4**: The recorded time series of the used variables after imputation and transformations.

#### **Section S1.3: Forecasting with EDM**

We forecasted the simulated species dynamics with empirical dynamic modeling (EDM, Ye *et al.* 2015; Ye & Sugihara 2016). EDM is a nonparametric and nonlinear modeling approach in which estimations and predictions are made based on when the considered system was in a similar state at other times (with the state of a system being determined by the states of its components, e.g. by the species abundances). In other words, the computations are based on how the system behaved when it was in similar positions in the state space, regardless of whether these positions happened recently or not. For the state space reconstruction (SSR), EDM makes use of Takens’ theorem which asserts that the time series of a state variable contains imprints of other variables that interacted with it (Takens 1981). This means that variables needed for the SSR but that are not directly available can be accounted for by appropriately lagging recorded time series. The SSR is thus done by using both lagged and non-lagged time series. The number of time series needed for the SSR is defined as the embedding dimension E.

##### **Section S1.3.1: Forecasting** **simulated time series**

As the simulations consisted of single species time series, for the forecasts we used simplex EDM (which uses a single time series for the SSR). In simplex EDM, the forecasting of a specific time point is done by averaging how the system evolved in the $E+1$ system states that are most similar to the state based on which a prediction is being made.

We trained the forecast models using the simulated and sub-sampled time series. We did the forecast using the optimal $E$ determined for each time series. The highest value for the embedding dimension that we tested was 8, chosen because the highest value for $\tau$ (i.e. the lag used in creating the simulations, see Section S1.1) was 7.67 (i.e. $\tau=7.67$, $r=0.3$, $r\tau=2.3$). The forecasts were always eight days ahead, meaning that the time points forecasted ahead depended on the sampling interval and ranged from 16 (0.5 days sampling interval) to one (eight days sampling interval). To evaluate the forecast models, we forecasted the same 25 time points for every simulated time series of a given length. For this we used evaluation time series created in the same way as the training time series but without the addition of noise. As a measure of forecast error, we computed the root mean square error (RMSE) of the model-based predictions. We standardized the computed RMSE values by dividing them with the standard deviation of the time series used for the forecasts. The expected value of the standardized RMSE for forecasts based on the average abundance (i.e. the baseline model) is one (i.e. $RMSE_{baseline}=1$), which means that forecasts that achieve RMSE values below one perform better than the baseline model, regardless of the used carrying capacity. Further, this is equivalent to saying that they show greater forecast skill than the baseline model, because the corresponding skill score ($SS$) is calculated as $SS=1-\frac{RMSE_{forecasts}}{RMSE_{baseline}}=1- RMSE_{forecasts}$.

##### **Section S1.3.2: Forecasting recorded field time series**

We forecasted the abundances of the phyto- and zooplankton classes (henceforth referred to as targets). In addition to these targets, we used the mean epilimnion temperature, the mean mixed layer depth, the mean mixed layer irradiance, and the concentrations of ammonium, phosphate, and nitrate as predictors.

We carried out the forecasts using the EDM technique multiview embedding (Ye *et al.* 2015; Ye & Sugihara 2016). The multiview embedding approach is an implementation of EDM that is capable of dealing with high-dimensional systems (Ye & Sugihara 2016). It constructs all possible E-dimensional SSRs based on the available time series and their lags, ranks them based on in-sample forecast skill and then selects the K best ones to compute an ensemble forecast. Because this method considers all possible SSRs, multiview embedding reduces the dimensionality of the system and thus allows the use of small values of E. This is crucial as the number of possible SSRs in high-dimensional systems quickly increases with E, such that it may become computationally infeasible to use large values for it.

Here, we set the embedding dimension to E=2 as we have previously shown that this method produces reliable results already with such a small value (Daugaard *et al.* 2022). We used a maximal lag of 3, i.e., we used the time series unlagged, lagged by 1 time point and lagged by 2 time points. Further, for each target group we used 40 values for K logarithmically spaced between 1 and the three fourths of the number of time points of the used dataset. For each target we then kept the model that resulted in the lowest forecast error in terms of RMSE. Note that because we carried out the forecasts with standardized time series, the resulting RMSE values are already standardized.

For all the datasets we did 12-days ahead forecasts, matching the lowest sampling frequency. This meant that the number of time points (i.e. steps) forecasted ahead depended on the sampling frequency (Table S1). With the reduced sampling frequency datasets, we forecasted target abundances using leave-one-out cross-validation. To reduce computational load, for the complete daily dataset and the datasets containing a reduced number of time points we carried out the forecasts using *k*-fold cross-validation. For this, we divided the time series into training data (in-sample data) and evaluation data (out-of-sample data, set to 21 time points). This meant that number of folds (i.e. *k*) depended on the time series length (Table S1).

**Table S1**: The field datasets used and their characteristics as well as details about their forecasting. For the forecasting, either a leave-one-out cross-validation or a k-fold cross-validation was performed, as described in the main text. For the k-fold cross-validation the number of time points in the evaluation data was fixed, meaning that the number of folds varied accordingly (k = number of time points in time series / number of time points in evaluation time series). The days forecasted ahead was fixed to 12 days for all datasets, meaning that the number of steps forecasted ahead depended on the sampling frequency (steps-ahead = product of days-ahead forecasted and the sampling frequency).

| **Dataset** | **Sampling frequency (samplings/day)** | **Proportion of time points** | **Number of time points** | **Cross-validation method** | **Time points in evaluation data** | **Number of folds (*k*)** | **Steps-ahead forecasted** | **Days-ahead forecasted** |
| --- | --- | --- | --- | --- | --- | --- | --- | --- |
| **Complete** | 1/1 | 12/12 | 994 | *k*-fold | 21 | 47 | 12 | 12 |
| **Reduced proportion of time points** | 1/1 | 9/12 | 745 | *k*-fold | 21 | 35 | 12 | 12 |
|  | 1/1 | 6/12 | 497 | *k*-fold | 21 | 23 | 12 | 12 |
|  | 1/1 | 3/12 | 248 | *k*-fold | 21 | 12 | 12 | 12 |
|  | 1/1 | 2/12 | 165 | *k*-fold | 21 | 8 | 12 | 12 |
|  | 1/1 | 1/12 | 82 | *k*-fold | 21 | 4 | 12 | 12 |
| **Reduced sampling frequency** | 1/1 | 1/12 | 82 | Leave-one-out | 1 | 82 | 12 | 12 |
|  | 1/3 | 1/12 | 82 | Leave-one-out | 1 | 82 | 4 | 12 |
|  | 1/6 | 1/12 | 82 | Leave-one-out | 1 | 82 | 2 | 12 |
|  | 1/12 | 1/12 | 82 | Leave-one-out | 1 | 82 | 1 | 12 |

#### **Section S1.4: Estimation of number and strength of interactions**

We determined which state variables were causally linked using a test of causation based on Convergent Cross Mapping (CCM, Sugihara *et al.* 2012) and the recommendations of Deyle et al (2016). We extended the methodology with a stringent convergence test that compares CCM skill between two variables using 20% and 50% of the data. This is done with 100 random subsets of the data. If the CCM skill is larger when 50% of the data is used in at least 95% of subsets, then we considered the test to be passed. This convergence test has previously been used by Merz *et al*. (2023). In this way we determined which variables interacted with each of the targets and the number of interactions of the target. In the case of the reduced data, we did this in each of the datasets separately. Subsequently, to have a single estimate per target at each sampling frequency and time series length, we averaged over the respective datasets to estimate the mean number of interactions.

To estimate the interaction strengths between causally linked state variables (based on the CCM analysis) we used Smap EDM (Deyle *et al.* 2016). This method calculates the interaction time series between interacting variables by iteratively constructing the interaction Jacobian $J_{ij}=\partial T_{i}\left( t+\tau\right)/\partial I_{j}\left( t \right)$at each time point, with $T_{i}$ and $I_{j}$ being different target and interactor variables, $t$ the time and $\tau$ the time step. Similar to the forecasting, this is done by utilizing the information of how the system reacted when it was in a comparable state at other time points. In addition, the nonlinearity parameter $\theta$ assigns bigger weights to more similar system states, i.e., it weighs the contribution of system states based on their proximity in the state space.

Here we used an extension of this method called regularized Smap EDM (Cenci *et al.* 2019), which can handle process noise by introducing a penalization function and penalization parameter $\lambda$. More specifically, we used the elastic net regularization which has an additional parameter $\alpha$ that determines the shape of the penalization function. We set to $\alpha=0.9$ and used target-specific values for $\theta$ and $\lambda$ which we determined with a grid search ($\lambda$: 30 values logarithmically spaced between 0 and 1; $\theta\in\left\{ 0.01, 0.1, 0.3, 0.5, 0.75, 1, 1.5, 2, 3, 4, 5, 6, 7, 8, 9 \right\}$).

### **Section S2: Additional figures and tables**

#### **Section S2.1: Forecast error as a function of sampling design and functional traits**

Regarding the simulation study, figure S5 shows that while bigger measurement errors resulted in larger forecast errors, neither it nor the length of the time series length (i.e. the number of time points) changed the positive correlation between the species growth rate and the optimal sampling frequency (i.e. the one the yielded the lowest forecast error).

Table S2 reports the regression (and ANOVA) tables for when the forecast error of the targets was used as the response variable in various analyses with the explanatory variables: sampling frequency, proportion of time points, target maximum net growth rates and target median body size. Table S3 then reports the association between these growth rates and body sizes. As can be seen we found a positive correlation between maximum net growth rates and body sizes, which is the opposite of what is usually the case (e.g. Bonner 2015). There could be a number of explanations for this, of which we briefly mention a few. First, in this study we considered a relatively narrow body size gradient (see e.g. Figure S6), which could mean that it is not broad enough to capture the negative relation between growth rates and body size. A second possibility is that smaller species do indeed have smaller higher growth rates, but because they might also be eaten more quickly than larger species their maximum net growth rate might effectively be smaller. This would mean that there is a mismatch between the estimated maximum net growth rates and true intrinsic growth rates. A third possible explanation is that because we did not work on a species level but on a grouped level this grouping somehow inverted this relation. Regardless of the reason of the positive correlation, this does not particularly affect the results or their interpretation. For instance, as expected from hypotheses and literature maximum net growth rates were positively correlated with forecast error, and yet growth rates were not a good criterion to select sampling frequencies. This is independent of the relation between growth rates and body sizes.

Figure S6 depicts the relations between the target body sizes and the respective forecast errors and interaction estimates.

#### **Section S2.2: Interaction estimates as a function of sampling design and functional traits**

Table S4 reports the regression (and ANOVA) tables for when the estimated number of interactions of the targets was used as the response variable in various analyses with the explanatory variables: sampling frequency, proportion of time points, target maximum net growth rates and target median body size.

Table S5 reports the regression (and ANOVA) tables for when the estimated mean interaction strength of the targets was used as the response variable in various analyses with the explanatory variables: sampling frequency, proportion of time points, target maximum net growth rates and target median body size.

Figure S7 depicts the random intercepts and random slopes estimated for the targets in the regression between the number of interactions (response) and the proportion of time points used (explanatory variable). As can be seen, whether we estimated a target to have more or fewer interactions when we used an increasingly higher proportion of time points in the estimation was target specific, as we estimated both negative and positive slopes.

Figure S8 illustrates that using fewer time points in the estimation led to a decrease in the variance of the number of interactions estimated across targets. In other words, with decreasing sample size the estimated number of interactions converged to a similar value regardless of target.

#### **Section S2.3: Influence of sampling design on the detection of correlations between variables**

Table S6 report the regression results for the relation between the forecast error and the interaction estimates of the targets, as well as for the relation between the interaction estimates (i.e. number and average interaction strength). For each dataset (defined by the sampling frequency used or the proportion of time points included) the regressions were fitted separately. Figure S9 illustrates this table graphically.

Tables S7 reports the same as Table S6, but only for the case in which we did 1 day ahead forecasts (instead 12 days). We did this only for the complete daily data. Figure S10 illustrates this table graphically.

#### **Section S2.4: Robustness analyses**

##### **Section S2.4.1: Random Forest forecast**

The results reported in the main text are based on the described EDM forecasts. Additionally, to investigate whether the main result (i.e. the relation between sampling frequency and forecast error) depended on the chosen forecasting method, we also forecasted the plankton abundances by using Random Forest. For this, we used the transformed time series (see main text) and the R-package “randomForest” (Liaw & Wiener 2002). Just as for EDM, we forecasted 12 days ahead by using the time series sub-sampled at different sampling frequencies and we allowed the predictors (which were the same as for the EDM forecasts) to lagged by using a maximal lag of 3, i.e., we used the time series unlagged, lagged by 1 time point and lagged by 2 time points.

Mirroring the EDM results, we found a clear negative relation between the sampling frequency and the abundance forecast errors across all plankton groups (Table S8 and Figure S11). In fact, the best Random Forest-based forecasts were achieved at the highest sampling frequency for all targets, thus confirming our results (Figure S11b).

##### **Section S2.4.2: Controlling for the covered time window across sampling frequencies**

Time series that are based on different sampling frequencies will inevitably differ in either the number of time points that they include or in the time window (i.e. the absolute time) that they cover (or, of course, they can differ in both regards, but crucially both things cannot be controlled at the same time). Because the number of time points impacts analyses and forecasts (see Figures 2 and 3 in the main text), in the main text we prioritized controlling for number of time points rather than for the time window. Further, and as described in the main text, we employed a sub-sampling scheme (i.e. based on multiple time series per sampling frequency) and a forecasting approach (i.e. the leave-one-out forecasts) designed to at least partially control for the different time windows for time series of different sampling frequencies.

However, it remains possible that the different time windows did have an effect on the results. Further, by consulting the literature we observed that the usual approach to investigate sampling frequency effects is to control that all time series cover the same amount of absolute time. In other words, either explicitly or implicitly the usual approach is to allow the time series recorded at higher frequencies to have more time points.

To investigate whether our results also hold when we follow the traditional approach and thus when we control for the time window rather than for the number of time points, we carried out additional forecasts. For every target (i.e. plankton group) we used the longest time series at the various sampling frequencies to forecast its abundance over the same time period (the time series included 82, 163, 325 and 973 time points respectively for the sampling intervals of once every 12 days, every six days, every 3 days and every day). We again did leave-one-out forecasts based on the EDM approach, but modified the approach slightly such that for every time series we only forecasted time points that were included at every sampling frequency. Other than that, we carried out the EDM forecasts as described in Section S1.3.

Matching our findings presented in the main text, we found a negative relation between sampling frequency and forecast error and we found that we achieved the best abundance forecasts at the highest sampling frequency for all but two target (Table S8 and Figure S12).

##### **Section S2.4.3: Forecasting 24 days ahead**

Corresponding to the longest considered sampling interval, in the main text we carried out and analyzed 12-days ahead forecasts. Here, we extend this to 24-days ahead forecasts to investigate whether it is still possible to make skillful forecasts at this time horizon, and, if yes, whether we still find the same relation between sampling frequency and forecast error as in the 12-days ahead case. Other than the different number of days forecasted ahead, the EDM forecast model specifics were the same as before.

We found weak evidence for the same qualitative relation between sampling frequency and forecast error when we doubled the number of days forecasted ahead, although the relation was clearer for the phytoplankton groups (Table S8, Figure S13). However, at this time horizon especially at the lower sampling frequencies the standardized forecast errors were often close to one or larger than one, indicating that these forecasts were not better than simply taking the average of the time series. This might explain why we found only weak evidence for the mentioned relation. Further, because of the generally high RMSE values, determining the optimal sampling frequency when the forecasts are not better than the baseline model (i.e. the average value) is not sensible. Thus, Figure S13b, which shows the optimal sampling frequency for 24-days ahead forecasts, is only included for completeness and is not further discussed.

In conclusion, the 24-days ahead forecasts showed that the forecast horizon was reached at 24 days (or earlier) more often at the lower sampling frequencies, while at the higher sampling frequencies skillful forecast remained possible for more plankton groups.

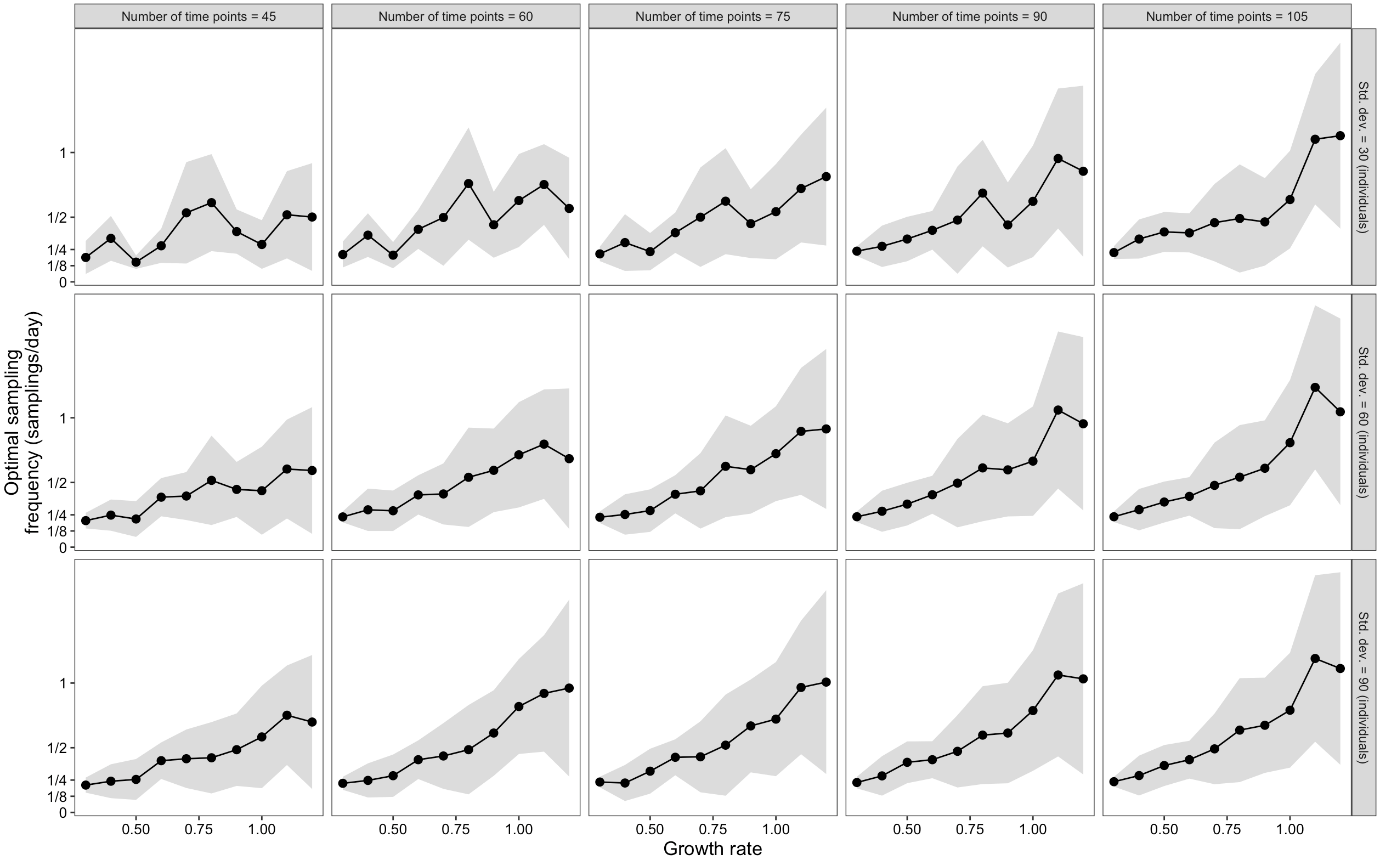

**Figure S5**: The relation growth rate and optimal sampling frequency in the simulations, as a function of number of time points in the time series and the amount of noise added (as specified in the subpanels). The results are shown as connected black dots (mean values across repeated simulations) and shaded regions (standard deviations).

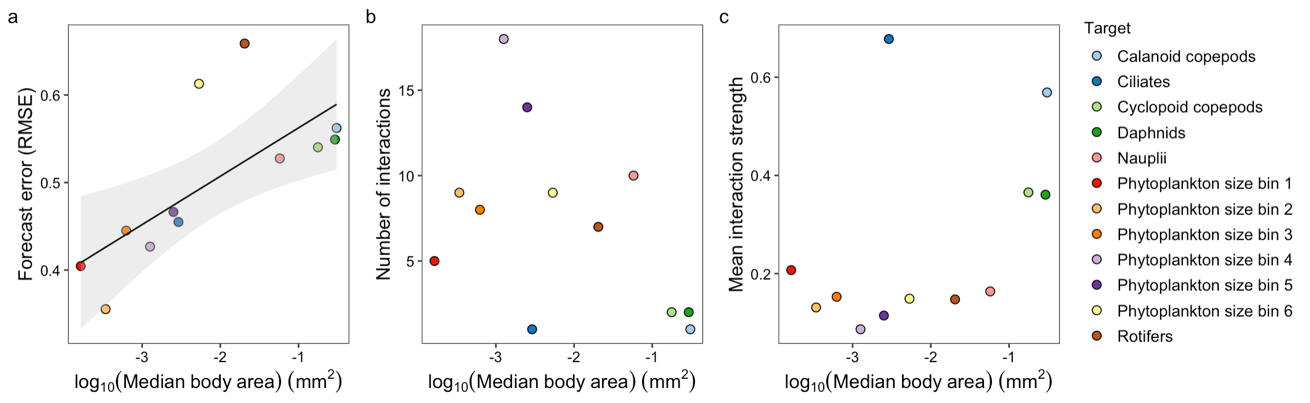

**Figure S6**: The relations between plankton class average body size and (**a**) the forecast error of the abundance of the class, (**b**) the estimated number of interactions of the class and (**c**) the mean interaction strength of the class. The dots are colored by target and the black lines and the shaded regions show the fixed effects results of the respective regressions and the corresponding 95% confidence intervals (if a significant relation was found).

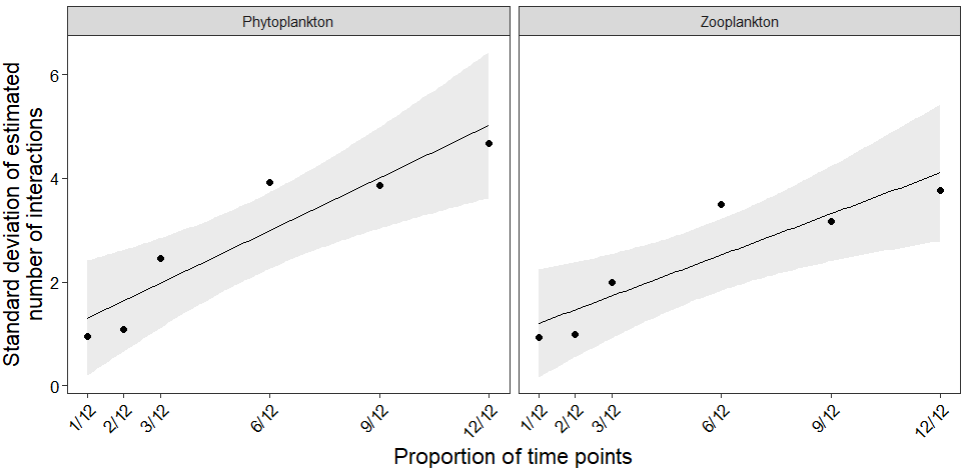

**Figure S8**: The standard deviation of the estimated number of interactions across all targets for a certain proportion of time points used, separately for the phyto- and zooplankton. The dots represent the estimated values, while the solid lines and the shaded regions show the results of a simple linear regression (estimated slope and corresponding 95% confidence interval).

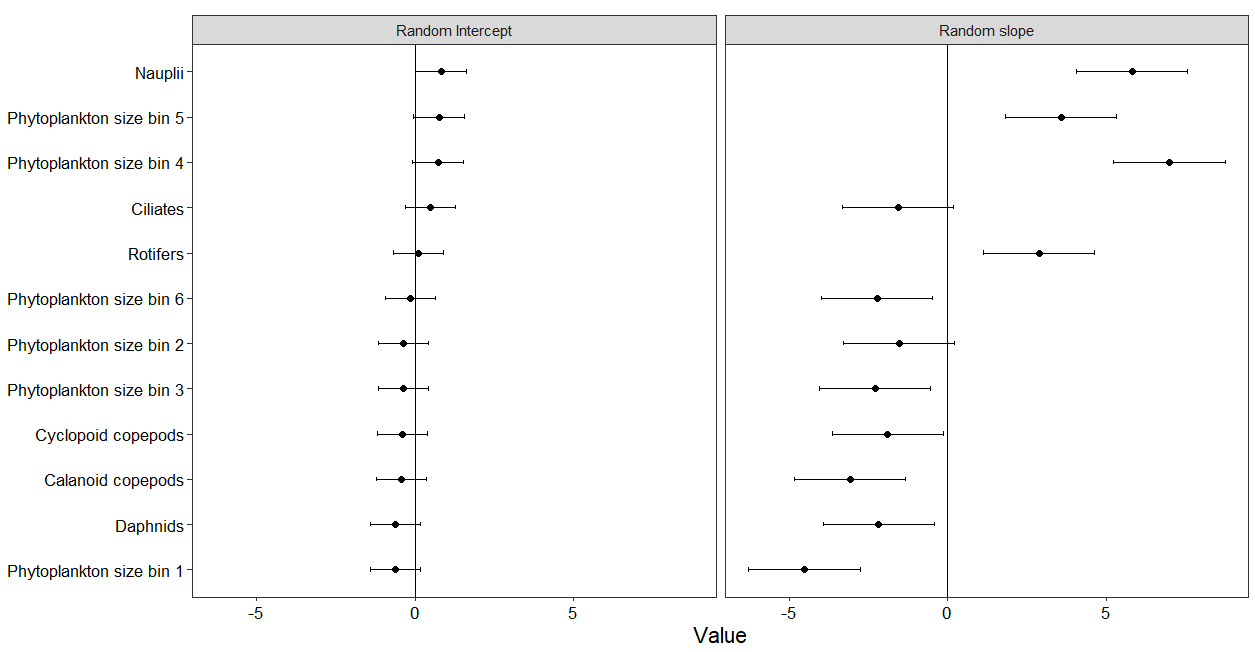

**Figure S7**: The estimated random intercepts and random slopes of the plankton classes for the relation between the proportion of time points used and the estimated number of interactions of the classes (ordered by decreasing random intercept). Dots represent the estimated values, while the error bars show the corresponding 95% confidence intervals.

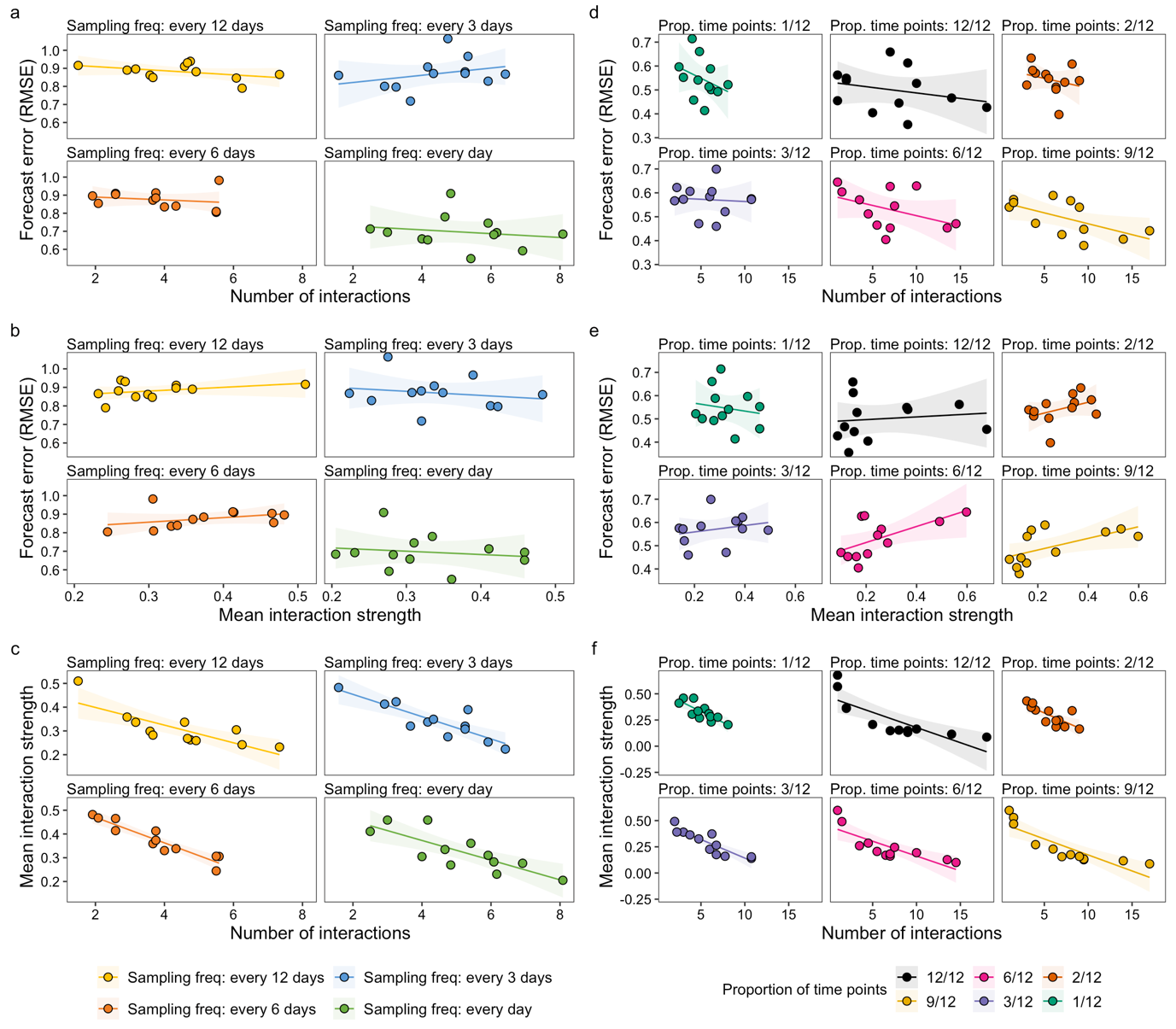

**Figure S9**: The relation between the number of interactions and the forecast error (**a** & **d**), between the mean interaction strength and the forecast error (**b** & **e**) and the relation between the number of interactions and the mean interaction strength (**c** & **f**). The left column (panels **a**, **b** and **c**) shows the relations for the datasets with reduced sampling frequency. The right column (panels **d**, **e** and **f**) shows the relations for the datasets with reduced number of time points. Each dot represents a plankton target and is colored based on the dataset used. The lines and shaded regions are colored by dataset used and represent the fixed effects results of the respective regressions (slope and 95% confidence intervals).

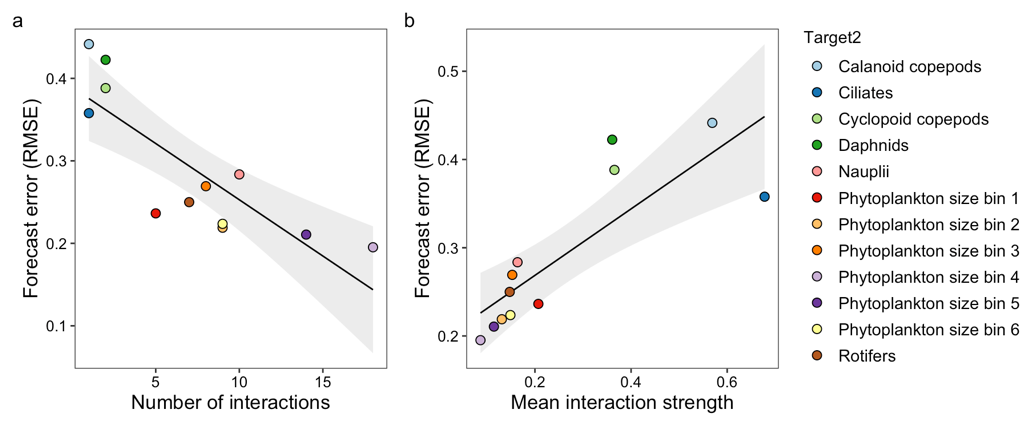

**Figure S10**: Figure showing the relation between estimated interactions and forecasting error for the complete daily data for when the number of steps forecasted ahead and the number of days forecasted ahead were both one. (a) The relation between the estimated number of interactions and the forecast error. (b)The relation between the estimated mean interaction strength and the forecast error. The dots are colored by target and the solid lines, and the shaded regions are the results of the respective regressions (slopes and 95% confidence intervals).

**Figure S11**: Random forest forecasts. (a) The relation between abundance forecast error (field data, random forest forecasts) and the sampling frequency. The dots are colored by target and the lines are the fixed effects of the corresponding regressions (shaded regions: 95% CI). (b) The relation between growth rate and the optimal sampling frequency for abundance forecasting. Connected black dots (mean values) and the shaded region (standard deviations) show the results for the simulations, with parameter values as in Figure 2a. The colored points show the relation for the targets in the field data (random forest forecast).

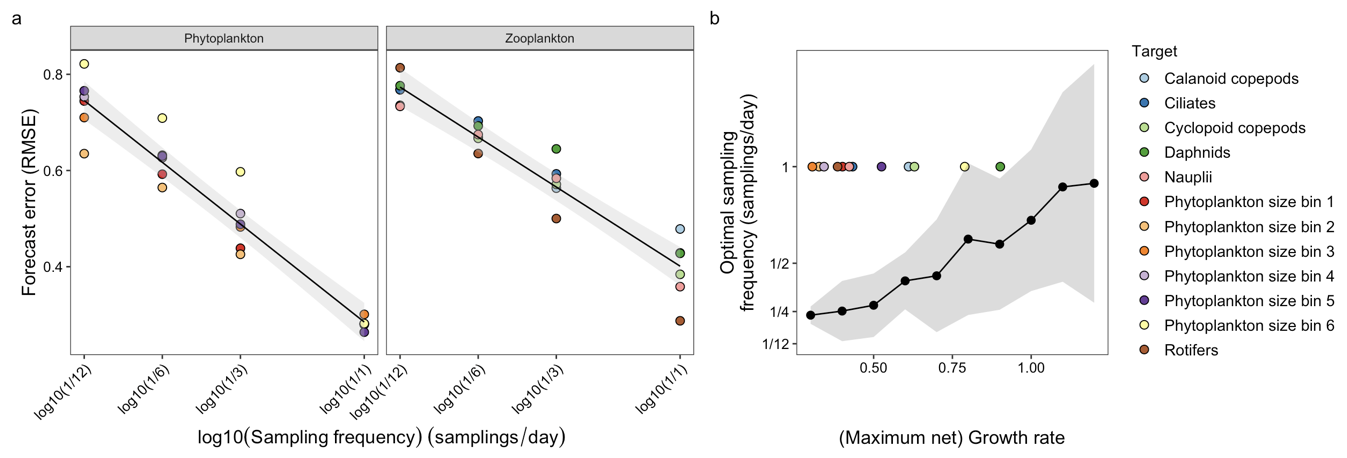

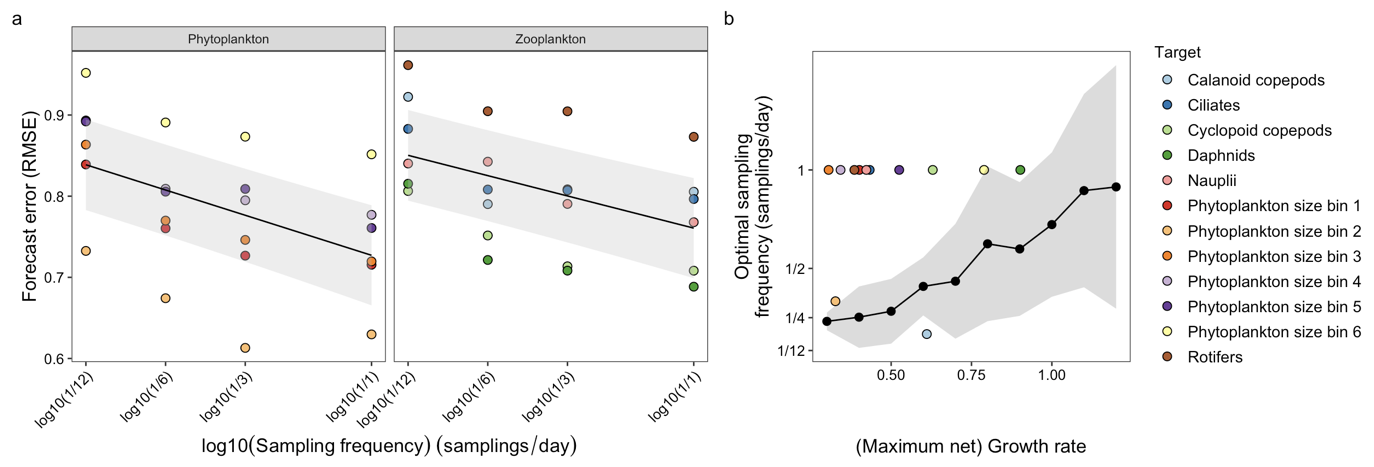

**Figure S12**: Controlled time window forecasts. (a) The relation between abundance forecast error (field data, time window controlled) and the sampling frequency. The dots are colored by target and the lines are the fixed effects of the corresponding regressions (shaded regions: 95% CI). (b) The relation between growth rate and the optimal sampling frequency for abundance forecasting. Connected black dots (mean values) and the shaded region (standard deviations) show the results for the simulations, with parameter values as in Figure 2a. The colored points show the relation for the targets in the field data (controlled time window).

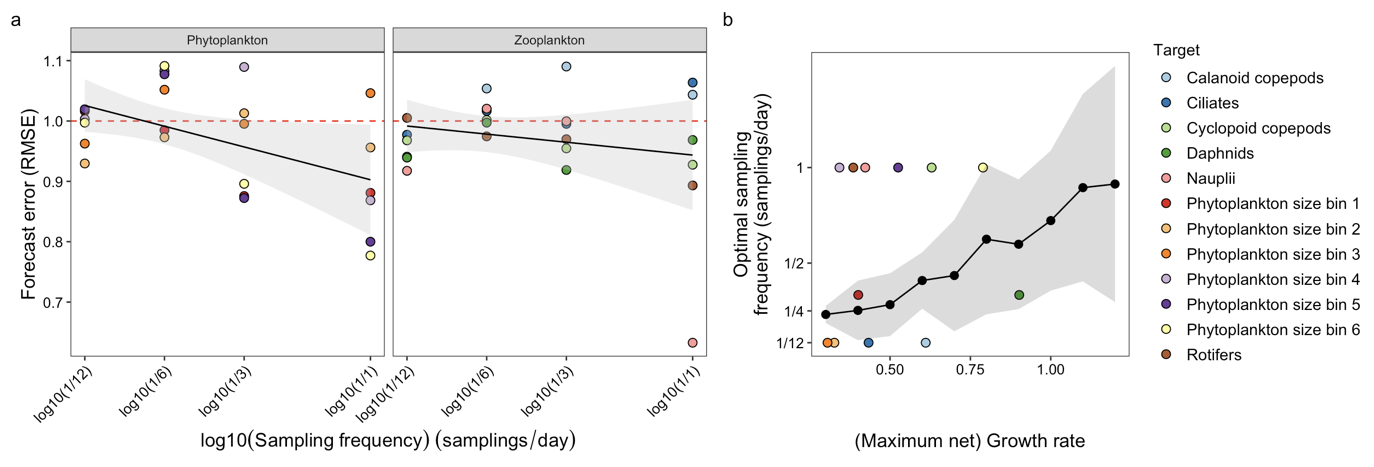

**Figure S13**: 24 days ahead forecasts. (a) The relation between abundance forecast error (field data, 24 days ahead) and the sampling frequency. The dots are colored by target and the lines are the fixed effects of the corresponding regressions (shaded regions: 95% CI). The horizontal line (red and dashed) has an intercept of 1, indicating the expected RMSE value for forecasts done by using the respective average abundance of the target time series. (b) The relation between growth rate and the optimal sampling frequency for abundance forecasting. Connected black dots (mean values) and the shaded region (standard deviations) show the results for the simulations, with parameter values as in Figure 2a. The colored points show the relation for the targets in the field data (24 days ahead).

**Table S2**: Forecast error analyses. The table reports the regression (and ANOVA Type III) results for when the forecast error was the response variable, and different explanatory variables were used in separate analyses (as specified in the column “Main covariate”). Note that when the explanatory variable was either the sampling frequency or the proportion of time points used then we fitted a mixed effects model with additionally also an interaction between the main covariate and the plankton group. Therefore, for completeness we also report the estimated random effects and the ANOVA table. Median area = Median body size. Net growth rate = maximum net growth rate.

| **Main covariate** | **Covariate** | **Type** | **Estimate** | **DF** | **SE** | **Test** | **Value** | ***p*-value** |
| --- | --- | --- | --- | --- | --- | --- | --- | --- |
| **Sampling frequency** | Intercept | Fix. eff. | 0.925 | 10 | 0.02 | *t* | 45.626 | <0.0001 |
|  | Sampling frequency | Fix. eff. | -0.259 | 10 | 0.046 | *t* | -5.669 | 0.0002 |
|  | Zooplankton | Fix. eff. | -0.021 | 10 | 0.029 | *t* | -0.747 | 0.4722 |
|  | Interaction | Fix. eff. | 0.095 | 10 | 0.065 | *t* | 1.466 | 0.1734 |
|  | Random intercept | Std. Dev. | 0.033 | - | - | - | - | - |
|  | Random slope | Std. Dev. | 0.087 | - | - | - | - | - |
|  | Sampling frequency | Sum of sq. | 0.109 | 1, 10 | - | *F* | 42.914 | <0.0001 |
|  | Plankton | Sum of sq. | 0.001 | 1, 10 | - | *F* | 0.558 | 0.4722 |
|  | Interaction | Sum of sq. | 0.005 | 1, 10 | - | *F* | 2.149 | 0.1734 |
| **Proportion of time points** | Intercept | Fix. eff. | 0.528 | 10 | 0.021 | *t* | 24.835 | <0.0001 |
|  | Time points prop. | Fix. eff. | -0.087 | 10 | 0.041 | *t* | -2.141 | 0.0580 |
|  | Zooplankton | Fix. eff. | 0.069 | 10 | 0.03 | *t* | 2.281 | 0.0457 |
|  | Interaction | Fix. eff. | 0.037 | 10 | 0.058 | *t* | 0.647 | 0.5323 |
|  | Random intercept | Std. Dev. | 0.042 | - | - | - | - | - |
|  | Random slope | Std. Dev. | 0.084 | - | - | - | - | - |
|  | Time points prop. | Sum of sq. | 0.011 | 1, 10 | - | *F* | 5.668 | 0.0386 |
|  | Plankton | Sum of sq. | 0.01 | 1, 10 | - | *F* | 5.204 | 0.0457 |
|  | Interaction | Sum of sq. | 0.001 | 1, 10 | - | *F* | 0.418 | 0.5323 |
| **Net growth rate** | Intercept | Fix. eff. | 0.609 | 10 | 0.054 | *t* | 11.345 | <0.0001 |
|  | log10(net growth rate) | Fix. eff. | 0.337 | 10 | 0.152 | *t* | 2.218 | 0.0509 |
| **Median area** | Intercept | Fix. eff. | 0.618 | 10 | 0.041 | *t* | 15.048 | <0.0001 |
|  | log10(median area) | Fix. eff. | 0.055 | 10 | 0.017 | *t* | 3.226 | 0.0091 |

**Table S3**: Regression table showing the relation between the maximum net growth rate of the targets and their mean body size (i.e. median area) on the log_10_-scale.

| **Response** | **Covariate** | **Type** | **Estimate** | **DF** | **SE** | **Test** | **Value** | ***p*-value** |
| --- | --- | --- | --- | --- | --- | --- | --- | --- |
| **log10(median area)** | Intercept | Fix. eff. | -0.463 | 10 | 0.620 | *t* | -0.748 | 0.4717 |
|  | log10(net growth rate) | Fix. eff. | 5.168 | 10 | 1.755 | *t* | 2.945 | 0.0147 |

**Table S4**: Number of interactions analyses. Similar to Table S2, but with the estimated number of interactions of the targets as the response variable. Note that, as explained in the main text when the explanatory variable was either the maximum net growth rate or the median area (i.e. body size), then the estimate number of interactions was a count variable. Therefore, we fitted a Poisson regression in these two cases. Median area = Median body size. Net growth rate = maximum net growth rate.

| **Main covariate** | **Covariate** | **Type** | **Estimate** | **DF** | **SE** | **Test** | **Value** | ***p*-value** |
| --- | --- | --- | --- | --- | --- | --- | --- | --- |
| **Sampling frequency** | Intercept | Fix. eff. | 4.172 | 10 | 0.604 | *t* | 6.902 | <0.0001 |
|  | Sampling frequency | Fix. eff. | 2.251 | 11 | 0.817 | *t* | 2.755 | 0.0190 |
|  | Zooplankton | Fix. eff. | -0.340 | 10 | 0.855 | *t* | -0.398 | 0.6991 |
|  | Interaction | Fix. eff. | -2.273 | 11 | 1.155 | *t* | -1.967 | 0.0753 |
|  | Random intercept | Std. Dev. | 1.286 | - | - | - | - | - |
|  | Random slope | Std. Dev. | 1.459 | - | - | - | - | - |
|  | Sampling frequency | Sum of sq. | 3.623 | 1, 10.8 | - | *F* | 3.723 | 0.0803 |
|  | Plankton | Sum of sq. | 0.154 | 1, 10 | - | *F* | 0.158 | 0.6991 |
|  | Interaction | Sum of sq. | 3.766 | 1, 10.8 | - | *F* | 3.869 | 0.0753 |
| **Proportion of time points** | Intercept | Fix. eff. | 6.499 | 10 | 0.425 | *t* | 15.305 | <0.0001 |
|  | Time points prop. | Fix. eff. | 4.694 | 10 | 1.746 | *t* | 2.689 | 0.0227 |
|  | Zooplankton | Fix. eff. | -2.386 | 10 | 0.601 | *t* | -3.973 | 0.0026 |
|  | Interaction | Fix. eff. | -4.834 | 10 | 2.469 | *t* | -1.958 | 0.0787 |
|  | Random intercept | Std. Dev. | 0.710 | - | - | - | - | - |
|  | Random slope | Std. Dev. | 4.058 | - | - | - | - | - |
|  | Time points prop. | Sum of sq. | 4.014 | 1, 10 | - | *F* | 3.404 | 0.0948 |
|  | Plankton | Sum of sq. | 18.619 | 1, 10 | - | *F* | 15.789 | 0.0026 |
|  | Interaction | Sum of sq. | 4.522 | 1, 10 | - | *F* | 3.835 | 0.0787 |
| **Net growth rate** | Intercept | Fix. eff. | 1.190 | 10 | 0.623 | *t* | 1.911 | 0.0851 |
|  | log10(net growth rate) | Fix. eff. | -2.265 | 10 | 1.607 | *t* | -1.410 | 0.1890 |
| **Median area** | Intercept | Fix. eff. | 1.229 | 10 | 0.527 | *t* | 2.334 | 0.0418 |
|  | log10(median area) | Fix. eff. | -0.320 | 10 | 0.195 | *t* | -1.641 | 0.1318 |

**Table S5**: Mean interaction strength analyses. Similar to Table S2 and Table S3, but with the estimated mean interaction strengths of the targets as the response variable. Median area = Median body size. Net growth rate = maximum net growth rate.

| **Main covariate** | **Covariate** | **Type** | **Estimate** | **DF** | **SE** | **Test** | **Value** | ***p*-value** |
| --- | --- | --- | --- | --- | --- | --- | --- | --- |
| **Sampling frequency** | Intercept | Fix. eff. | 0.335 | 10 | 0.034 | *t* | 9.906 | <0.0001 |
|  | Sampling frequency | Fix. eff. | -0.055 | 10 | 0.052 | *t* | -1.075 | 0.3076 |
|  | Zooplankton | Fix. eff. | 0.014 | 10 | 0.048 | *t* | 0.297 | 0.7722 |
|  | Interaction | Fix. eff. | 0.084 | 10 | 0.073 | *t* | 1.158 | 0.2739 |
|  | Random intercept | Std. Dev. | 0.072 | - | - | - | - | - |
|  | Random slope | Std. Dev. | 0.101 | - | - | - | - | - |
|  | Sampling frequency | Sum of sq. | 0 | 1, 10 | - | *F* | 0.131 | 0.7245 |
|  | Plankton | Sum of sq. | 0 | 1, 10 | - | *F* | 0.088 | 0.7722 |
|  | Interaction | Sum of sq. | 0.004 | 1, 10 | - | *F* | 1.340 | 0.2739 |
| **Proportion of time points** | Intercept | Fix. eff. | 0.26 | 10 | 0.036 | *t* | 7.242 | <0.0001 |
|  | Time points prop. | Fix. eff. | -0.147 | 10 | 0.080 | *t* | -1.848 | 0.0943 |
|  | Zooplankton | Fix. eff. | 0.095 | 10 | 0.051 | *t* | 1.861 | 0.0923 |
|  | Interaction | Fix. eff. | 0.167 | 10 | 0.113 | *t* | 1.485 | 0.1685 |
|  | Random intercept | Std. Dev. | 0.075 | - | - | - | - | - |
|  | Random slope | Std. Dev. | 0.177 | - | - | - | - | - |
|  | Time points prop. | Sum of sq. | 0.006 | 1, 10 | - | *F* | 1.276 | 0.2850 |
|  | Plankton | Sum of sq. | 0.015 | 1, 10 | - | *F* | 3.465 | 0.0923 |
|  | Interaction | Sum of sq. | 0.01 | 1, 10 | - | *F* | 2.204 | 0.1685 |
| **Net growth rate** | Intercept | Fix. eff. | 0.401 | 10 | 0.132 | *t* | 3.038 | 0.0125 |
|  | log10(net growth rate) | Fix. eff. | 0.438 | 10 | 0.374 | *t* | 1.172 | 0.2686 |
| **Median area** | Intercept | Fix. eff. | 0.418 | 10 | 0.113 | *t* | 3.707 | 0.0041 |
|  | log10(median area) | Fix. eff. | 0.074 | 10 | 0.047 | *t* | 1.577 | 0.1460 |

**Table S6**: Regression tables for the relation between the forecast error and the interaction estimates, as well as between the interaction estimates themselves (i.e. number and mean strength of interactions). For each dataset (defined by either the sampling frequency or the proportion of time points used) the regressions were carried out separately.

| **Response** | **Dataset** | **Covariate** | **Estimate** | **DF** | **SE** | ***t*-value** | ***p*-value** | **95% CI** | |
| --- | --- | --- | --- | --- | --- | --- | --- | --- | --- |
| **Forecast error (RMSE)** | Sampling every day | Intercept | 0.750 | 10 | 0.094 | 7.942 | <0.0001 | 0.539 | 0.960 |
|  |  | Num. interactions | -0.011 | 10 | 0.018 | -0.598 | 0.5635 | -0.050 | 0.029 |
|  | Sampling every 3 days | Intercept | 0.782 | 10 | 0.087 | 8.957 | <0.0001 | 0.587 | 0.976 |
|  |  | Num. interactions | 0.020 | 10 | 0.019 | 1.052 | 0.3173 | -0.022 | 0.062 |
|  | Sampling every 6 days | Intercept | 0.906 | 10 | 0.047 | 19.084 | <0.0001 | 0.800 | 1.011 |
|  |  | Num. interactions | -0.008 | 10 | 0.012 | -0.669 | 0.5186 | -0.035 | 0.019 |
|  | Sampling every 12 days | Intercept | 0.932 | 10 | 0.035 | 26.774 | <0.0001 | 0.855 | 1.010 |
|  |  | Num. interactions | -0.011 | 10 | 0.007 | -1.545 | 0.1533 | -0.028 | 0.005 |
| **Forecast error (RMSE)** | Sampling every day | Intercept | 0.754 | 10 | 0.115 | 6.560 | <0.0001 | 0.498 | 1.010 |
|  |  | Mean int. strength | -0.178 | 10 | 0.343 | -0.518 | 0.6156 | -0.942 | 0.587 |
|  | Sampling every 3 days | Intercept | 0.945 | 10 | 0.125 | 7.536 | <0.0001 | 0.666 | 1.225 |
|  |  | Mean int. strength | -0.221 | 10 | 0.360 | -0.616 | 0.5520 | -1.023 | 0.580 |
|  | Sampling every 6 days | Intercept | 0.783 | 10 | 0.076 | 10.313 | <0.0001 | 0.614 | 0.952 |
|  |  | Mean int. strength | 0.246 | 10 | 0.199 | 1.236 | 0.2446 | -0.197 | 0.689 |
|  | Sampling every 12 days | Intercept | 0.819 | 10 | 0.052 | 15.718 | <0.0001 | 0.703 | 0.936 |
|  |  | Mean int. strength | 0.202 | 10 | 0.165 | 1.224 | 0.2492 | -0.166 | 0.570 |
| **Mean interaction strength** | Sampling every day | Intercept | 0.539 | 10 | 0.052 | 10.416 | <0.0001 | 0.423 | 0.654 |
|  |  | Num. interactions | -0.041 | 10 | 0.010 | -4.308 | 0.0015 | -0.063 | -0.020 |
|  | Sampling every 3 days | Intercept | 0.548 | 10 | 0.041 | 13.491 | <0.0001 | 0.457 | 0.638 |
|  |  | Num. interactions | -0.047 | 10 | 0.009 | -5.311 | 0.0003 | -0.067 | -0.027 |
|  | Sampling every 6 days | Intercept | 0.578 | 10 | 0.024 | 23.721 | <0.0001 | 0.523 | 0.632 |
|  |  | Num. interactions | -0.054 | 10 | 0.006 | -8.770 | <0.0001 | -0.067 | -0.040 |
|  | Sampling every 12 days | Intercept | 0.473 | 10 | 0.042 | 11.398 | <0.0001 | 0.381 | 0.566 |
|  |  | Num. interactions | -0.037 | 10 | 0.009 | -4.222 | 0.0018 | -0.057 | -0.018 |
| **Forecast error (RMSE)** | Prop. time points: 1/12 | Intercept | 0.640 | 10 | 0.083 | 7.699 | <0.0001 | 0.455 | 0.825 |
|  |  | Num. interactions | -0.018 | 10 | 0.015 | -1.175 | 0.2671 | -0.053 | 0.016 |
|  | Prop. time points: 2/12 | Intercept | 0.589 | 10 | 0.057 | 10.295 | <0.0001 | 0.462 | 0.717 |
|  |  | Num. interactions | -0.008 | 10 | 0.010 | -0.853 | 0.4134 | -0.029 | 0.013 |
|  | Prop. time points: 3/12 | Intercept | 0.582 | 10 | 0.046 | 12.774 | <0.0001 | 0.481 | 0.684 |
|  |  | Num. interactions | -0.002 | 10 | 0.007 | -0.264 | 0.7971 | -0.017 | 0.014 |
|  | Prop. time points: 6/12 | Intercept | 0.590 | 10 | 0.044 | 13.293 | <0.0001 | 0.491 | 0.689 |
|  |  | Num. interactions | -0.009 | 10 | 0.006 | -1.533 | 0.1564 | -0.021 | 0.004 |
|  | Prop. time points: 9/12 | Intercept | 0.562 | 10 | 0.032 | 17.373 | <0.0001 | 0.490 | 0.634 |
|  |  | Num. interactions | -0.009 | 10 | 0.004 | -2.466 | 0.0333 | -0.017 | -0.001 |
|  | Prop. time points: 12/12 | Intercept | 0.533 | 10 | 0.045 | 11.872 | <0.0001 | 0.433 | 0.633 |
|  |  | Num. interactions | -0.005 | 10 | 0.005 | -0.898 | 0.3903 | -0.016 | 0.007 |
| **Forecast error (RMSE)** | Prop. time points: 1/12 | Intercept | 0.601 | 10 | 0.106 | 5.680 | 0.0002 | 0.366 | 0.837 |
|  |  | Mean int. strength | -0.170 | 10 | 0.316 | -0.537 | 0.6028 | -0.875 | 0.535 |
|  | Prop. time points: 2/12 | Intercept | 0.465 | 10 | 0.057 | 8.153 | <0.0001 | 0.338 | 0.592 |
|  |  | Mean int. strength | 0.267 | 10 | 0.187 | 1.427 | 0.1841 | -0.150 | 0.685 |
|  | Prop. time points: 3/12 | Intercept | 0.532 | 10 | 0.053 | 10.095 | <0.0001 | 0.414 | 0.649 |
|  |  | Mean int. strength | 0.138 | 10 | 0.170 | 0.813 | 0.4354 | -0.241 | 0.518 |
|  | Prop. time points: 6/12 | Intercept | 0.445 | 10 | 0.040 | 11.100 | <0.0001 | 0.355 | 0.534 |
|  |  | Mean int. strength | 0.345 | 10 | 0.139 | 2.488 | 0.0321 | 0.036 | 0.654 |
|  | Prop. time points: 9/12 | Intercept | 0.431 | 10 | 0.032 | 13.408 | <0.0001 | 0.360 | 0.503 |
|  |  | Mean int. strength | 0.251 | 10 | 0.105 | 2.377 | 0.0388 | 0.016 | 0.486 |
|  | Prop. time points: 12/12 | Intercept | 0.485 | 10 | 0.047 | 10.418 | <0.0001 | 0.381 | 0.589 |
|  |  | Mean int. strength | 0.058 | 10 | 0.146 | 0.399 | 0.6983 | -0.267 | 0.383 |
| **Mean interaction strength** | Prop. time points: 1/12 | Intercept | 0.539 | 10 | 0.052 | 10.416 | <0.0001 | 0.423 | 0.654 |
|  |  | Num. interactions | -0.041 | 10 | 0.010 | -4.308 | 0.0015 | -0.063 | -0.020 |
|  | Prop. time points: 2/12 | Intercept | 0.498 | 10 | 0.060 | 8.346 | <0.0001 | 0.365 | 0.632 |
|  |  | Num. interactions | -0.036 | 10 | 0.010 | -3.648 | 0.0045 | -0.058 | -0.014 |
|  | Prop. time points: 3/12 | Intercept | 0.497 | 10 | 0.037 | 13.367 | <0.0001 | 0.414 | 0.580 |
|  |  | Num. interactions | -0.035 | 10 | 0.006 | -6.233 | <0.0001 | -0.048 | -0.023 |
|  | Prop. time points: 6/12 | Intercept | 0.445 | 10 | 0.053 | 8.466 | <0.0001 | 0.328 | 0.562 |
|  |  | Num. interactions | -0.028 | 10 | 0.007 | -4.273 | 0.0016 | -0.043 | -0.014 |
|  | Prop. time points: 9/12 | Intercept | 0.478 | 10 | 0.050 | 9.498 | <0.0001 | 0.366 | 0.590 |
|  |  | Num. interactions | -0.031 | 10 | 0.006 | -5.308 | 0.0003 | -0.043 | -0.018 |
|  | Prop. time points: 12/12 | Intercept | 0.468 | 10 | 0.060 | 7.837 | <0.0001 | 0.335 | 0.601 |
|  |  | Num. interactions | -0.029 | 10 | 0.007 | -4.270 | 0.0016 | -0.044 | -0.014 |

**Table S7**: the same as Table S6, but only considering the case in which 1-day ahead forecasts were done (instead of 12-days ahead) and the entire time series were used (i.e. all time points and the highest sampling frequency).

| **Response** | **Covariate** | **Estimate** | **DF** | | **SE** | ***t*-value** | ***p*-value** | **95% CI** | |
| --- | --- | --- | --- | --- | --- | --- | --- | --- | --- |
| **Forecast error (RMSE)** | Intercept | 0.389 | 10 | 0.025 | | 15.335 | <0.0001 | 0.333 | 0.446 |
|  | Num. interactions | -0.014 | 10 | 0.003 | | -4.731 | 0.0008 | -0.020 | -0.007 |
| **Forecast error (RMSE)** | Intercept | 0.193 | 10 | 0.026 | | 7.503 | <0.0001 | 0.136 | 0.251 |
|  | Mean int. strength | 0.377 | 10 | 0.081 | | 4.664 | 0.0009 | 0.197 | 0.557 |

**Table S8**: Robustness analyses. The table reports the regression (and ANOVA Type III) results for when the forecast error was the response variable, and different robustness analyses were carried out (as specified in the column “Robustness analysis”). Note that we fitted a mixed effects model with additionally also an interaction between the main covariate and the plankton group. Therefore, for completeness we also report the estimated random effects and the ANOVA table.

| **Robustness analysis** | **Covariate** | **Type** | **Estimate** | **DF** | **SE** | **Test** | **Value** | ***p*-value** |
| --- | --- | --- | --- | --- | --- | --- | --- | --- |
| **Random forest forecasts** | Intercept | Fix. eff. | 0.285 | 10 | 0.02 | *t* | 14.033 | <0.0001 |
|  | Sampl. freq | Fix. eff. | -0.427 | 10 | 0.026 | *t* | -16.157 | <0.0001 |
|  | Zooplankton | Fix. eff. | 0.116 | 10 | 0.029 | *t* | 4.042 | 0.0024 |
|  | Interaction | Fix. eff. | 0.082 | 10 | 0.037 | *t* | 2.189 | 0.0534 |
|  | Rand. int. | Std. Dev. | 0.044 | - | - | - | - | - |
|  | Rand. slope | Std. Dev. | 0.056 | - | - | - | - | - |
|  | Sampl. freq | Sum of sq. | 0.295 | 1, 10 | - | *F* | 426.86 | <0.0001 |
|  | Plankton | Sum of sq. | 0.011 | 1, 10 | - | *F* | 16.335 | 0.0024 |
|  | Interaction | Sum of sq. | 0.003 | 1, 10 | - | *F* | 4.794 | 0.0534 |
| **Time window control** | Intercept | Fix. eff. | 0.727 | 10 | 0.031 | *t* | 23.105 | <0.0001 |
|  | Sampl. freq | Fix. eff. | -0.103 | 29 | 0.014 | *t* | -7.37 | <0.0001 |
|  | Zooplankton | Fix. eff. | 0.033 | 10 | 0.045 | *t* | 0.753 | 0.4690 |
|  | Interaction | Fix. eff. | 0.020 | 29 | 0.02 | *t* | 1.022 | 0.3152 |
|  | Rand. int. | Std. Dev. | 0.073 | - | - | - | - | - |
|  | Rand. slope | Std. Dev. | 0.006 | - | - | - | - | - |
|  | Sampl. freq | Sum of sq. | 0.064 | 1, 29 | - | *F* | 88.369 | <0.0001 |
|  | Plankton | Sum of sq. | 0.0004 | 1, 10 | - | *F* | 0.566 | 0.4690 |
|  | Interaction | Sum of sq. | 0.001 | 1, 29 | - | *F* | 1.044 | 0.3152 |
| **24 days ahead forecasts** | Intercept | Fix. eff. | 0.903 | 10 | 0.047 | *t* | 19.318 | <0.0001 |
|  | Sampling frequency | Fix. eff. | -0.114 | 10 | 0.056 | *t* | -2.037 | 0.0677 |
|  | Zooplankton | Fix. eff. | 0.041 | 10 | 0.066 | *t* | 0.619 | 0.5497 |
|  | Interaction | Fix. eff. | 0.069 | 10 | 0.079 | *t* | 0.877 | 0.4002 |
|  | Random intercept | Std. Dev. | 0.1 | - | - | - | - | - |
|  | Random slope | Std. Dev. | 0.112 | - | - | - | - | - |
|  | Sampling frequency | Sum of sq. | 0.016 | 1, 10 | - | *F* | 4.017 | 0.0716 |
|  | Plankton | Sum of sq. | 0.002 | 1, 10 | - | *F* | 0.383 | 0.5497 |
|  | Interaction | Sum of sq. | 0.003 | 1, 10 | - | *F* | 0.769 | 0.4002 |

### **Section S4: Versions of R and the R-packages used**

**R version** 4.1.2 (2021-11-01)

**Base packages:** parallel, stats, graphics, grDevices, utils, datasets, methods, base

**Other packages**:

kableExtra 1.3.4,	odin 1.2.4,	lattice 0.20-45,

viridis 0.6.2,	viridisLite 0.4.0,	forecast 8.16,

truncnorm 1.0-8,	RColorBrewer 1.1-2,	merTools 0.5.2,

arm 1.12-2,	MASS 7.3-54,	lmerTest 3.1-3,

lme4 1.1-27.1,	Matrix 1.4-0,	ggpubr 0.4.0,

slider 0.2.2,	ggfortify 0.4.13,	elasticnet 1.3,

lars 1.2,	here 1.0.1,	GGally 2.1.2,

lubridate 1.8.0,	doParallel 1.0.16,	iterators 1.0.13,

foreach 1.5.1,	rEDM 1.13.1,	pracma 2.3.6,

data.table 1.14.2,	patchwork 1.1.1,	forcats 0.5.,

stringr 1.4.0,	dplyr 1.0.7,	purr 0.3.4,

readr 2.1.1,	tidyr 1.1.4,	tibble 3.1.6,

ggplot2 3.4.0,	tidyverse 1.3.1,	spatstat 2.3-3,

spatstat.linnet 2.3-2,	spatstat.core 2.4-0,	rpart 4.1-15,

nlme 3.1-153,	spatstat.random 2.1-0,	spatstat.geom 2.3-2,

spatstat.data 2.1-2,	randomForest 4.6-14,
